# Local and global Cdc42 GEFs for fission yeast cell polarity are coordinated by microtubules and the Tea1/Tea4/Pom1 axis

**DOI:** 10.1101/211912

**Authors:** Ye Dee Tay, Marcin Leda, Andrew B. Goryachev, Kenneth E. Sawin

## Abstract

The conserved Rho-family GTPase Cdc42 plays a central role in eukaryotic cell polarity. The rod-shaped fission yeast *Schizosaccharomyces pombe* has two Cdc42 guanine-nucleotide exchange factors (GEFs), Scd1 and Gef1, but little is known about how they are coordinated in polarized growth. Although the microtubule cytoskeleton is normally not required for polarity maintenance in fission yeast, here we show that when *scdl* function is compromised, disruption of microtubules or the polarity landmark proteins Tea1, Tea4, or Pom1 leads to isotropic rather than polarized growth. Surprisingly, this isotropic growth is due to spatially inappropriate activity of Gef1, which is a cytosolic protein rather than a membrane-associated protein at cell tips like Scd1. Microtubules and the Tea1/Tea4/Pom1 axis counteract inappropriate Gef1 activity by regulating the localization of the Cdc42 GTPase-activating protein Rga4. Our results thus demonstrate coordination of “local” (Scd1) and “global” (Gef1) Cdc42 GEFs via microtubules and microtubule-dependent polarity landmarks.

## INTRODUCTION

Cell polarity is fundamental to eukaryotic cells and essential for a range of functions, such as migration and/or directional growth, intracellular transport, cell signaling, asymmetric cell division, and tissue organization (Campanale et al., 2017; Mayor and Etienne-Manneville, 2016; Rodriguez-Boulan and Macara, 2014; Schelski and Bradke, 2017; St Johnston and Ahringer, 2010). Processes involved in cell polarization include the generation of spatial cues (intrinsic or extrinsic) for polarity site selection, recruitment of specific proteins to regions of plasma membrane, and reorganization of the actin and microtubule cytoskeleton and of intracellular trafficking. The Rho-family GTPase Cdc42 has important roles in many of these processes (Chiou et al., 2017; Etienne-Manneville, 2004, 2013; Hall, 2012; Harris and Tepass, 2010; Martin and Arkowitz, 2014; Perez and Rincon, 2010). Like other small GTPases, Cdc42 binds effector proteins when in its active, GTP-bound, state. Thus, control of Cdc42 activity by GTPase activating proteins (GAPs) and guanine-nucleotide exchange factors (GEFs) is a critical feature of polarity regulation (Bos et al., 2007; Cook et al., 2014; Hodge and Ridley, 2016; Moon and Zheng, 2003; Rossman et al., 2005).

Unicellular eukaryotes such as budding yeast *Saccharomyces cerevisiae* and fission yeast *Schizosaccharomyces pombe* are excellent models for studying Cdc42-dependent cell polarity, due to their simple geometries and reduced complexity relative to metazoan cells (Chiou et al., 2017; Martin and Arkowitz, 2014). Budding yeast are ovoid and form a single bud once per cell cycle, while fission yeast are rod-shaped and grow at their tips. In recent years, work in budding yeast has led to key insights into the mechanism(s) by which a stable, self-organized polarity cluster based on Cdc42-GTP can emerge on the plasma membrane to establish a presumptive bud site (Chiou et al., 2017; Goryachev and Leda, 2017; Woods and Lew, 2017). Cdc42 cluster formation depends on spontaneous symmetry-breaking via likely multiple converging positive feedback loops involving active Cdc42, Cdc42 effectors, such as the polarity scaffold protein Bem1, and the Cdc42 GEF, Cdc24.

The local enrichment of these factors via positive feedback can be sufficient for the establishment of cell polarity at a site designated by internal and/or external cues (Chiou et al., 2017; Goryachev and Leda, 2017; Woods and Lew, 2017).

While many of the components and mechanisms involved in budding yeast polarity are also conserved in fission yeast, there are also distinct differences. Scd1, the fission yeast ortholog of budding yeast Cdc24, is thought to have a similar role to Cdc24, functioning in a positive feedback loop to organize polarity clusters of active Cdc42 on the plasma membrane at cell tips, via interaction with Cdc42 effectors and scaffold protein Scd2 (Chang et al., 1999; Chang et al., 1994; Chiou et al., 2017; Endo et al., 2003). However, while Cdc24 is essential for viability, Scd1 is non-essential. In addition to Scd1, fission yeast has a second Cdc42 GEF, Gef1. Scd1 and Gef1 are thought to share an overlapping essential function, because single-deletion mutants of either gene *(scd1Δ* or *gef1Δ)* are viable, while the double-deletion mutant *(scd1Δ gef1Δ)* is lethal (Coll et al., 2003; Hirota et al., 2003).

Both Scd1 and Gef1 have been described to localize to the cell tips during interphase and to the cell mid-zone during cytokinesis (Coll et al., 2003; Das et al., 2009; Hirota et al., 2003) (Das et al., 2015; Kokkoris et al., 2014; Vjestica et al., 2013). However, phenotypes associated with both deletion and overexpression differ significantly between Scd1 and Gef1. Unlike rod-shaped wild-type cells, *scd1Δ* cells have a mostly round morphology (Chang et al., 1994) and lack detectable enrichment of Cdc42-GTP at cell tips (Kelly and Nurse, 2011; Tatebe et al., 2008). By contrast, *gef1Δ* cells have a largely wild-type morphology, albeit with mild defects in bipolar tip growth and septum formation (Coll et al., 2003). Similarly, Scd1 over-expression leads to no significant change in cell morphology, whereas Gef1 over-expression causes cells to become wider or rounder (Coll et al., 2003; Das et al., 2012). It is currently unclear how Scd1 and Gef1 activities are coordinated in the activation of Cdc42 at cell tips.

Another significant difference between fission yeast and budding yeast is that in fission yeast, interphase MTs make important contributions to cell polarity regulation (Martin and Arkowitz, 2014) (Chiou et al., 2017). This is not the case in budding yeast (Huffaker et al., 1988; Jacobs et al., 1988), and thus in this regard, fission yeast may be more similar to mammalian cells, in which MTs can interact directly or indirectly with multiple polarity regulators and also provide tracks for directed transport of vesicles and signaling molecules (Etienne-Manneville, 2013; Neukirchen and Bradke, 2011; Siegrist and Doe, 2007; Sugioka and Sawa, 2012).

Interphase MTs in fission yeast are nucleated from multiple intracellular sites and form ~3-5 bundles, each containing ~2-5 MTs, that extend along the long axis of the cell (Chang and Martin, 2009; Sawin and Tran, 2006). Landmark proteins such as Tea1 and Tea4 are continuously delivered to the cell tip via the plus ends of dynamic MTs (Martin et al., 2005; Mata and Nurse, 1997; Tatebe et al., 2005). The landmark proteins further recruit polarity factors such as the protein kinase Pom1, PP1 protein phosphatase Dis2, formin For3 and actin-associated protein Bud6, (Alvarez-Tabares et al., 2007; Bahler and Pringle, 1998; Glynn et al., 2001; Martin et al., 2005).

The importance of MTs in fission yeast cell polarity has been demonstrated by pharmacological inhibition (Sawin and Nurse, 1998; Sawin and Snaith, 2004) and by mutation of genes involved in microtubule biogenesis and function (Hirata et al., 1998; Radcliffe et al., 1998; Umesono et al., 1983; Vardy and Toda, 2000) (Anders et al., 2006; Samejima et al., 2005; Sawin et al., 2004). In general, two related polarity phenotypes are observed: mutations affecting MT nucleation and organization often lead to curved cells, while mutations affecting landmark proteins tend to lead to bent or branched cells, particularly after stress. Both of these phenotypes, however, contrast sharply with those associated with mutations in the Cdc42 polarity module, which lead to round-or wide-cell phenotypes (Chang et al., 1994; Kelly and Nurse, 2011; Miller and Johnson, 1994). Collectively, these findings have led to a general model in which MTs and MT-dependent landmark proteins play an important role in selecting sites for polarity establishment but are not required for polarity establishment *per se,* or for the maintenance of polarized growth (Chang and Martin, 2009; Sawin and Snaith, 2004). At the same time, these differences in phenotypes (i.e. mispositioned polarity vs. loss or impairment of polarity) highlight the limitations in our understanding of how MT and MT-dependent landmarks may contribute to regulation of Cdc42-dependent cell polarity.

Here we simultaneously address both the question of how the two fission yeast Cdc42 GEFs are coordinated in cell polarity regulation and how MTs and their effectors contribute to regulation of the core cell polarity machinery. Previous work has shown that polarized growth can be observed in interphase *scd1Δ* cells if interphase is extended by preventing cell cycle progression (Kelly and Nurse, 2011). Here we find, surprisingly, that polarized growth in *scd1Δ* cells absolutely requires interphase MTs; after MT disruption, *scd1Δ* cells grow isotropically, resulting in round cells. We further show that MTs promote polarized growth in *scd1* mutants via a pathway involving polarity proteins Tea1, Tea4, and Pom1 (the Tea1/Tea4/Pom1 “axis”), as well as Cdc42 GAP Rga4 (Das et al., 2007; Kokkoris et al., 2014; Tatebe et al., 2008). Remarkably, this pathway serves to counteract the activity of Gef1, which we find, contrary to the published literature (Das et al., 2015; Das et al., 2009; Kokkoris et al., 2014; Vjestica et al., 2013), to be a cytosolic “global” Cdc42 GEF, rather than a membrane-associated “local” GEF like Scd1. Our results reveal a previously unrecognized role of MTs and the Tea1/Tea4/Pom1 axis in the maintenance of cell polarity. We propose a model for fission yeast cell polarity regulation in which local and global Cdc42 GEFs are active in parallel but are regulated by different mechanisms. If not coordinated, these can impair rather than promote polarized growth.

## RESULTS

### Polarized growth of scd1Δ cells

Recently it was shown that *scd1Δ* cells arrested in G1/S-phase by hydroxyurea treatment have a polarized shape (Kelly and Nurse, 2011). This suggested that *scd1Δ* cells may normally be polarized, but because of their round shape, polarization can be observed unambiguously only during an extended interphase. To investigate polarization of *scd1Δ* without using hydroxyurea, we overexpressed the CDK-inhibitory kinase Wee1 *(adh13:wee1)* in *scd1Δ* cells that also expressed CRIB-3mCitrine, a reporter for active (GTP-bound) Cdc42 (Jaquenoud and Peter, 2000; Mutavchiev et al., 2016; Tatebe et al., 2008). Compared to *scd1Δ* cells, *adh13:wee1 scd1Δ* cells were clearly polarized, with an aspect ratio (length divided by width) similar to that of wild-type cells (Fig. 1A,B). Interestingly, in spite of this polarization, *adh13:wee1 scd1Δ* cells did not show any detectable CRIB-3mCitrine at cell tips (Fig. 1A), similar to previous observations in hydroxyurea-arrested cells (Kelly and Nurse, 2011).

**Figure 1.**
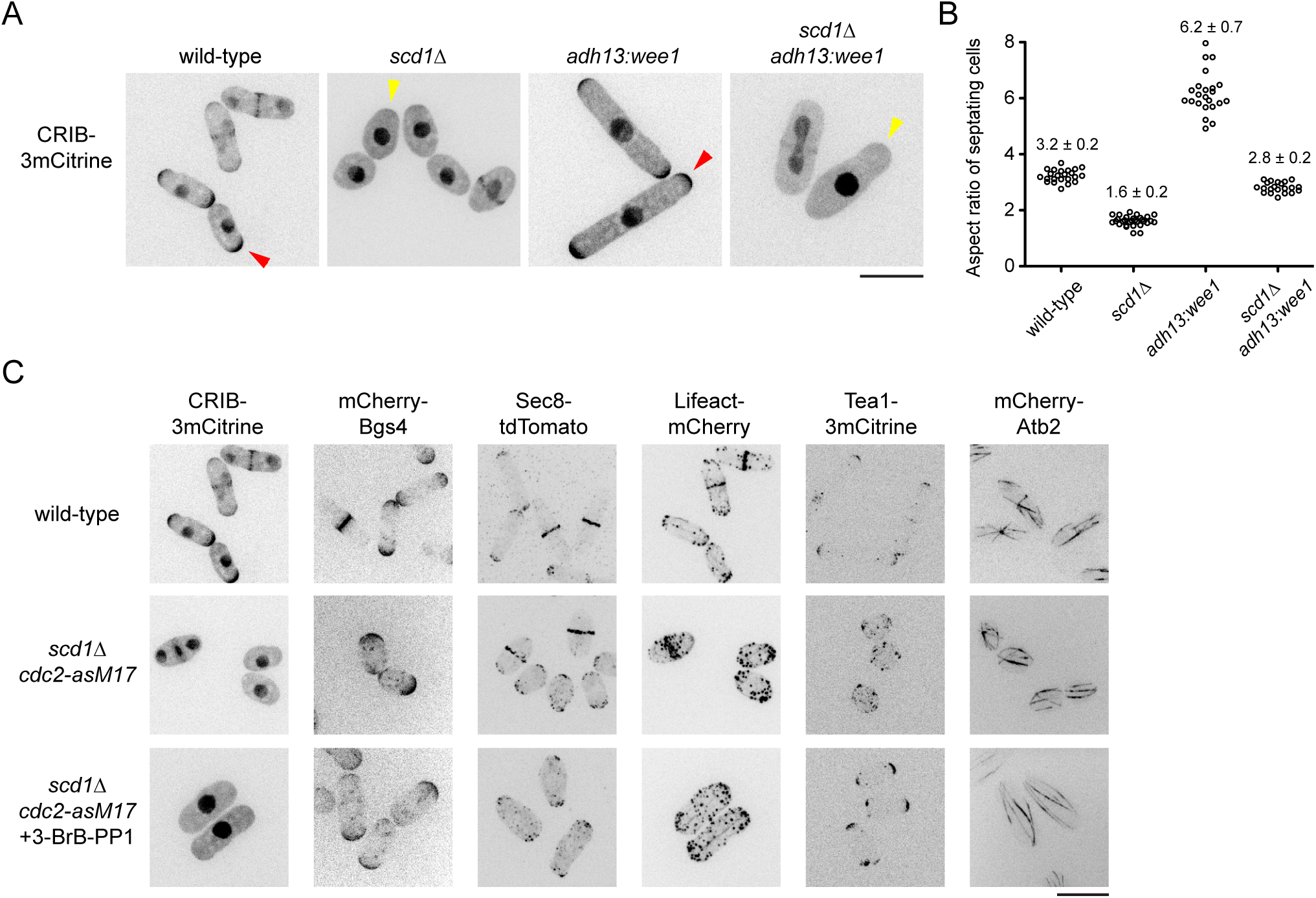
*scd1Δ* cells undergo polarized growth during extended interphase, but with nearly undetectable Cdc42-GTP at cell tips. **(A)** Cell morphology and distribution of Cdc42-GTP reporter CRIB-3mCitrine in cells of the indicated genotypes. Polarized shape of ***scd**1Δ* cells is clearly seen upon ***wee1*** overexpression. Presence and absence of CRIB at cell tips is indicated by red and yellow arrowheads, respectively. **(B)** Aspect ratio (cell length divided by cell width) of septating cells of the indicated genotypes, with mean and SD. Wild-type and ***scd**1Δ **adh13:wee1*** cells have similar ratios. For each strain, 21-30 cells were scored. **(C)** Localization of several different polarity-associated proteins in cells of the indicated genotypes. In bottom panels, cells were treated with 3-BrB-PP1 for 5 hr prior to imaging. Polarity markers other than CRIB are enriched at cell tips. Scale bars, 10 μm. See also Figure 1--figure supplement 1.

To characterize *scd1Δ* polarized growth in further detail, we used *cdc2-asM17* cells, which have a mutation in the ATP-binding pocket of Cdc2 and thus can be arrested in interphase by treatment with nucleotide-competitive analogs (Aoi et al., 2014; Bishop et al., 2000; Cipak et al., 2011). In addition to CRIB-3mCitrine, we imaged several different fluorescent-tagged cell polarity reporters in *scd1Δ cdc2-asM17* cells, including beta-glucan synthase Bgs4 (Cortes et al., 2005; Cortes et al., 2015), exocyst component Sec8 (Snaith et al., 2011; Wang et al., 2002), F-actin reporter Lifeact (Huang et al., 2012; Riedl et al., 2008), and polarity landmark Tea1 (Mata and Nurse, 1997) (Fig. 1C). After treatment with the nucleotide-competitive analog 3-BrB-PP1, *scd1Δ cdc2-asM17* cells were clearly polarized, and all polarity reporters localized to cell tips, with the exception of CRIB-3mCitrine, which, as in *scd1Δ adh13:wee1* cells, was not seen at cell tips. The absence of CRIB-3mCitrine from cell tips in *scd1Δ cdc2-asM17* was further confirmed by imaging additional reporters of the Scd1-Cdc42 polarity module (Fig. 1—figure supplement 1). In untreated *scd1Δ cdc2-asM17* cells, polarity reporters (excepting CRIB-3mCitrine) could also be seen at cell tips, although for some proteins (e.g. Lifeact, Tea1), the round cell shape made this more difficult to discern (Fig. 1C).

We conclude that *scd1Δ* cells can grow in a polarized manner, with nearly all of the hallmarks of normal polarized growth. Due to the increased width of *scd1Δ* cells, their polarized growth is most easily apparent when interphase is extended. In addition, *scd1Δ* polarized growth is not associated with detectable levels of the CRIB reporter at cell tips (Kelly and Nurse, 2011).

### Polarized growth in scd1 mutants depends on microtubules and on polarity landmark proteins Tea1 and Tea4

Failure to detect CRIB-3mCitrine at cell tips in *scd1Δ* cells prompted us to ask what other factors might be important for *scd1Δ* polarized growth. Although MTs are not required for polarized growth in wild-type *(scd1+)* cells ((Sawin and Snaith, 2004); see Introduction), we hypothesized that MTs might contribute specifically to polarized growth in *scd1Δ* cells.

We imaged mCherry-Bgs4 in *scd1Δ cdc2-asM17* cells during extended interphase after 3-BrB-PP1 treatment, both in the presence and absence of the MT-depolymerizing drug MBC (Fig. 2, Video 1). Inhibition of Cdc2-asM17 by 3-BrB-PP1 was particularly useful because it allowed imaging of cell growth for several hours without intervening cell division. In the absence of MBC, *scd1Δ cdc2-asM17* grew in a polarized manner, as did control *(scd1+) cdc2-asM17* cells in the presence of MBC. Strikingly, after addition of MBC to *scd1Δ cdc2-asM17* cells, Bgs4 no longer localized mainly to cell tips and instead formed transient, mobile patches on the plasma membrane (Fig. 2A). Accordingly, instead of growing in a polarized manner, these cells grew isotropically, becoming increasingly round over time (Fig. 2B,C). We conclude that MTs are critical for polarized growth in *scd1Δ* cells, but not in wild-type *(scd1+)* cells.

**Figure 2.**
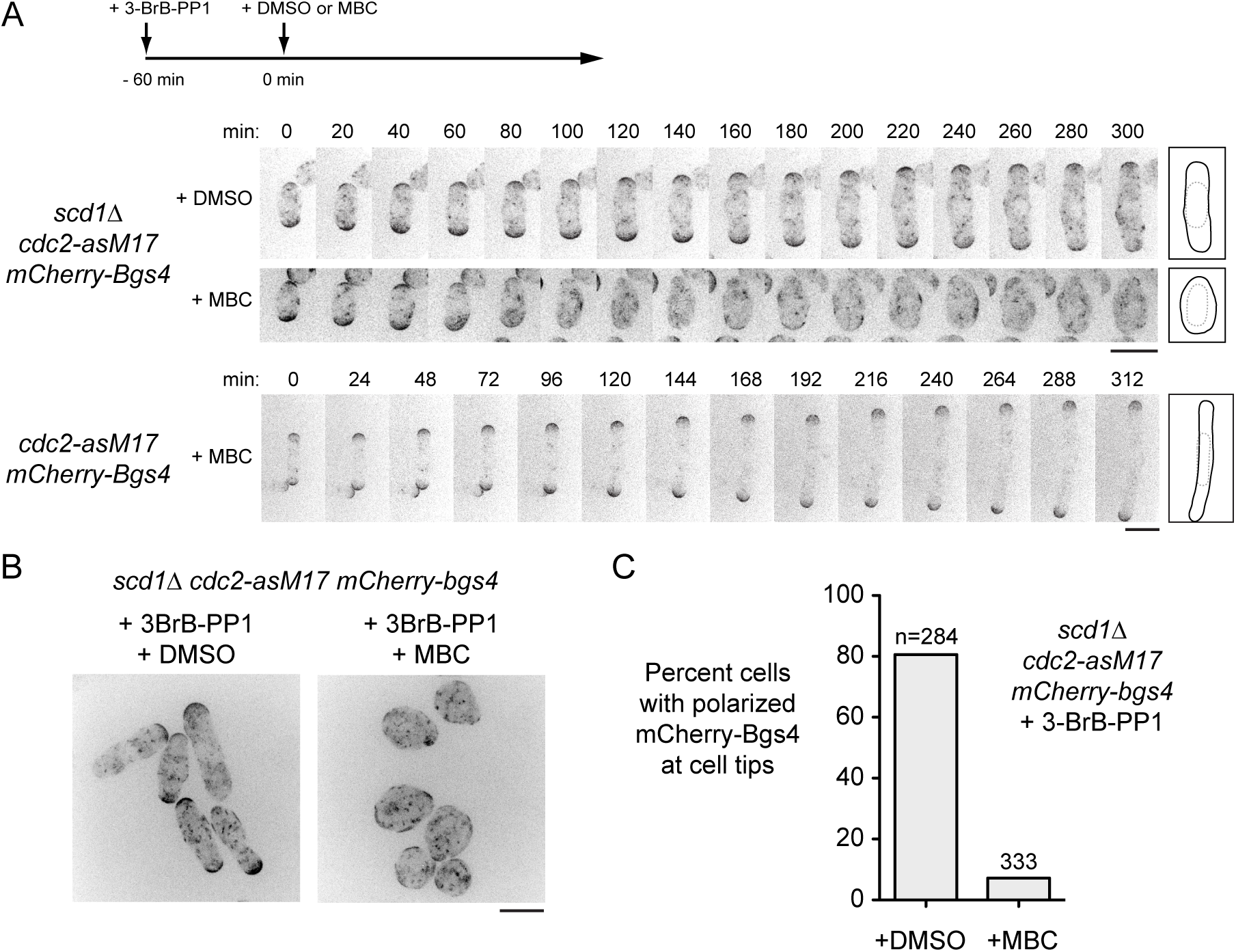
Microtubule depolymerization in *scd1Δ* cells leads to isotropic growth. **(A)** Time courses from movies showing mCherry-Bgs4 distribution and cell morphology in *scd1Δ cdc2-asM17 mCherry-bgs4* cells (top two rows) and control *cdc2-asM17 mCherry-bgs4* cells (bottom row) pre-treated with 3-BrB-PP1 at −60 min and then treated with either DMSO or MBC as indicated. Diagrams show cell outlines at beginning and end of movies; outlines were aligned slightly to account for limited cell movement. **(B)** Fields of cells as in A, after 3-BrB-PP1 pre-treatment and DMSO or MBC treatment for 300 min. **(C)** Percent cells containing polarized mCherry-Bgs4 during DMSO or MBC treatment (see Methods); n indicates total number of cells scored (polarized plus non-polarized). The difference between DMSO and MBC treatment was highly significant (p <0.0001; Fisher’s exact test). Scale bars, 10 pm. See also Video 2.

We hypothesized that MTs may contribute to polarized growth in *scd1Δ* cells via landmark proteins Tea1 and Tea4. Interestingly, and consistent with this hypothesis, *tea1Δ scd1Δ* double mutants are inviable (Papadaki et al., 2002). Therefore, to construct double mutants of *scd1* with *tea1Δ* and *tea4Δ,* we generated a strain in which expression of 3HA-tagged Scd1 is controlled by the weak, thiamine-repressible *nmt81* promoter (Basi et al., 1993) (Fig. 3—figure supplement 1). For simplicity, we will refer to the repressed *nmt81:3HA-scd1* allele as *scd1^low^.* Under repressing conditions, Scd1^low^ protein was not detected by Western blot, and, like *scd1Δ* cells, *scd1*^*low*^ cells had a round morphology and lacked detectable CRIB-3mCitrine at cell tips. We note, however, that other mutant phenotypes (see below) indicate that some biologically relevant, functional Scd1 is produced in these cells, albeit at very low levels.

**Figure 3.**
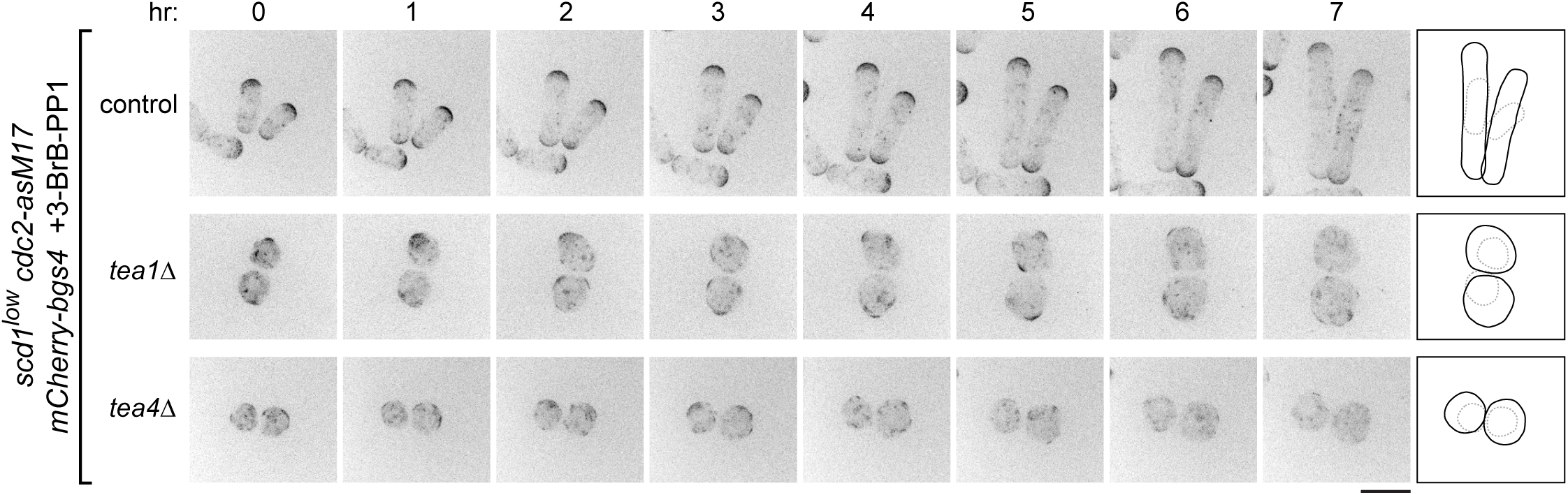
When *scd1* is expressed at very low levels, *tea1Δ* and *tea4A* cells grow isotropically. Time courses from movies showing mCherry-Bgs4 distribution and cell morphology in cells of the indicated genotypes (all are in ***scd1^low^ cdc2-asM17 mCherry-bgs4*** background). ***scd1*** expression was repressed with thiamine for 24 hr before imaging. 3-BrB-PP1 was added 30 minutes before imaging. Diagrams at right show cell outlines at beginning and end of movies; outlines were aligned slightly to account for limited cell movement. Note that during isotropic growth, transient “patches” of mCherry-Bgs4 appear at varied random positions on the cell surface, whereas during polarized growth, mCherry-Bgs4 remains at cell tips. Scale bar, 10 μm. See also Figure 3--figure supplement 1 and Video 2.

We next introduced *tea1Δ* and *tea4Δ* mutations into *scd1^low^ mCherry-Bgs4 cdc2-asM17* backgrounds. Under repressing conditions, *tea1A scd1^low^ mCherry-Bgs4 cdc2-asM17* and *tea4A scd1^low^ mCherry-Bgs4 cdc2-asM17* were viable but showed slightly increased frequency of cell death (see Methods). We repressed Scd1 expression for 24 h and then imaged cells after addition of 3-BrB-PP1 (Fig. 3; Video 2). In control cells, mCherry-Bgs4 remained highly polarized at cell tips after 3-BrB-PP1 addition, and cells grew in a polarized manner. By contrast, in *tea1Δ* and *tea4Δ* mutants, mCherry-Bgs4 was not polarized, even before 3-BrB-PP1 addition. Instead, in these cells mCherry-Bgs4 was present on internal membranes and on the plasma membrane as small, randomly-positioned patches, some of which were barely detectable. After 3-BrB-PP1 addition, *tea1A scd1^low^ mCherry-Bgs4 cdc2-asM17* and *tea4Δ scd1^low^ mCherry-Bgs4 cdc2-asM17* grew almost completely isotropically, with transient, mobile mCherry-Bgs4 patches (Fig. 3; Video 2; Fig. 4B). These results indicate that when Scd1 is expressed at very low levels, the absence of either Tea1 or Tea4 leads to near-complete loss of polarity, resulting in isotropic growth.

**Figure 4.**
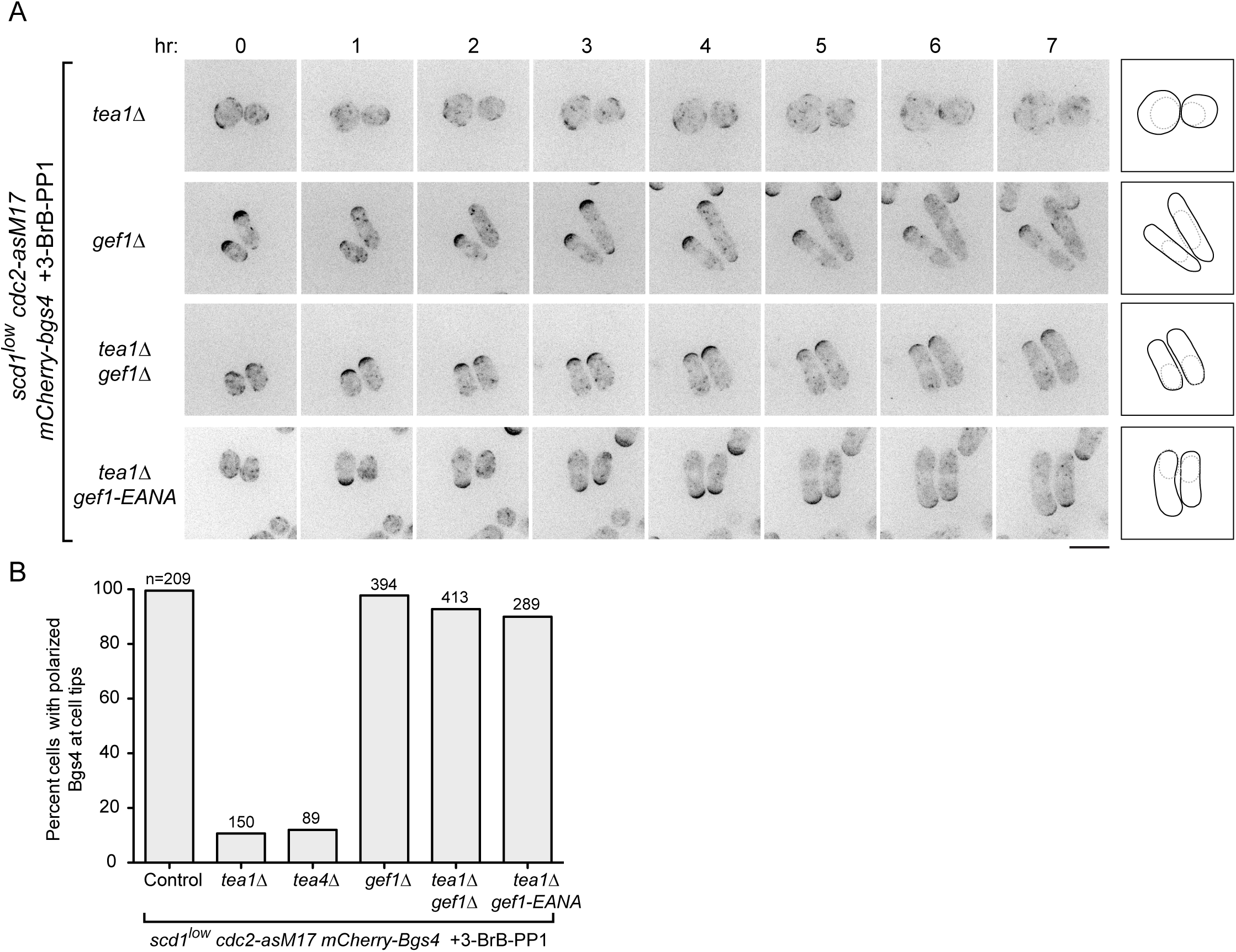
Loss of *gef1* function restores polarized growth to *scd1^low^ tea1Δ* cells. ***(A)*** Time courses from movies showing mCherry-Bgs4 distribution and cell morphology in cells of the indicated genotypes (all are in ***scd1^low^ cdc2-asM17 mCherry-bgs4*** background), at the indicated times after start of imaging. ***scd1*** expression was repressed with thiamine for 24 hr before imaging. 3-BrB-PP1 was added 30 minutes before imaging. Diagrams show cell outlines at beginning and end of movies; outlines were aligned slightly to account for limited cell movement. Note that newborn daughter cells often have less mCherry-Bgs4 at cell tips (e.g. in ***scd1^low^ cdc2-asM17 mCherry-bgs4 tea**1Δ **gef1-EANA*** cells). ***(B)*** Percent cells containing polarized mCherry-Bgs4 at cell tips, from movies of the type shown in Fig. 4A and Fig. 3; n indicates total number of cells scored (polarized plus non-polarized). mCherry-Bgs4 polarization was measured during the first four hours of imaging, when mCherry-Bgs4 signal remains strong. Pairwise differences relative to control were highly significant for all strains except ***gef**1Δ*, (p <0.0001; Fisher’s exact test, with correction for multiple comparisons). Scale bar, 10 μm. See also Figure 4--figure supplement 1 and Video 3.

### gef1 loss-of-function relieves the requirement for Tea1 and Tea4 in scd1^low^ polarized growth

We next tested whether Gef1 contributes to polarized growth when *scd1* function is compromised (Fig. 4). Like *tea1Δ scd1Δ* double mutants, *gef1Δ scd1Δ* double mutants are inviable (see Introduction). We therefore introduced *gef1Δ* into *scd1^low^ mCherry-Bgs4 cdc2-asM17* cells. Under repressing conditions, *gef1Δ scd1^low^ mCherry-Bgs4 cdc2-asM17* cells remained viable. Moreover, mCherry-Bgs4 was strongly enriched at cell tips both before and after 3-BrB-PP1 addition, and cells grew in a highly polarized manner (Fig. 4A,B; Video 3).

These results indicate that Gef1 is not required for polarized growth in *scd1*^*low*^ cells and therefore that the very low level of Scd1 expressed in *scd1*^*low*^ cells is sufficient for viability and polarized growth. This in turn raised the question of why Tea1 and Tea4 are required for polarized growth in *scd1*^*low*^ cells.

We hypothesized two possible roles for the Tea1/Tea4 system. The first possibility was that Tea1 and Tea4 might enhance the intrinsic ability of Scd1 to serve as a GEF when expressed at very low levels. The second possibility, which was motivated by the observation that Gef1 overexpression causes cell rounding (Coll et al., 2003; Das et al., 2012), was that rather than supporting Scd1 function directly, Tea1 and Tea4 may prevent or counteract any inappropriate function of Gef1, which would otherwise somehow interfere with the ability of low levels of Scd1 to promote polarized growth.

To distinguish between these possibilities, we introduced *gef1Δ* into *tea1Δ scd1^low^ mCherry-Bgs4 cdc2-asM17* cells and imaged cells after *scd1* repression and 3-BrB-PP1 addition. Remarkably, *gef1Δ* completely reversed the isotropic growth of *tea1Δ scd1^low^ mCherry-Bgs4 cdc2-asM17* cells, which now grew in a highly polarized manner, as seen both by cell shape and by mCherry-Bgs4 enrichment at cell tips (Fig. 4A,B; Video 3). This provides strong support for the second of the two possible roles proposed above.

In addition to a central catalytic Dbl homology (DH) domain, Gef1 contains an N-terminal region of unknown function and a C-terminal region that is proposed to contain a Bin/amphiphysin/Rvs (BAR) domain (Das et al., 2015). Because *gef1Δ* removes all of these domains, we wanted to determine whether the polarized growth seen in *gef1Δ tea1Δ scd1^low^ mCherry-Bgs4 cdc2-asM17* cells was specifically due to a loss of Gef1’s GEF activity. We therefore mutated conserved residues E318 and N505 in the Gef1 DH domain to generate a mutant *(E318A, N505A;* termed *gef1-EANA)* that, based on previous structural and *in vitro* biochemical analyses, should be folded properly but unable to bind Cdc42 (Aghazadeh et al., 1998; Rossman et al., 2005; Rossman et al., 2002a; Rossman et al., 2002b). Consistent with this, we found that *gef1-EANA* is a loss-of-function mutation, even though Gef1-EANA protein localized *in vivo* identically to wild-type Gef1 (Fig. 4—figure supplement 1). In further imaging experiments, we found that after *scd1* repression, *gef1-EANA tea1Δ scd1^low^ mCherry-Bgs4 cdc2-asM17* cells were polarized both before and after 3-BrB-PP1 addition (Fig. 4A,B; Video 3). This indicates that the reversal of isotropic growth seen in our experiments can be attributed specifically to the loss of Gef1 GEF activity, rather than to the absence of Gef1 protein more generally.

Collectively, these results suggest not only that Gef1 is not required for polarized growth in *scd1*^*low*^ cells but also that preventing or counteracting Gef1 activity is a prerequisite for polarized growth in *scd1*^*low*^ cells. According to this view, the main role of Tea1 (and Tea4) in promoting polarized growth in *scd1*^*low*^ cells is to prevent isotropic growth caused by inappropriate Gef1 activity; correspondingly, if Gef1 is not present, then Tea1 is no longer required for polarized growth.

### During unperturbed interphase, Gef1 is cytosolic rather than membrane-associated

How is Gef1 localized *in vivo* such that it can promote isotropic growth in *scd1*^*low*^cells? Initial characterization of Gef1 showed that it localized to the septum during cell division but did not have any specific localization during interphase (Coll et al., 2003; Hirota et al., 2003). However, it was later reported that Gef1 is also localized to cell tips during interphase (Das et al., 2015; Das et al., 2009; Vjestica et al., 2013). Because it was not obvious to us how cell tip-localized Gef1 would lead to isotropic growth, we reinvestigated Gef1 interphase localization.

Using several different fluorescent Gef1 fusion proteins, including previously published ones, we observed Gef1 at the septum during cell division, but we did not observe any specific localization of Gef1 during interphase, even with sensitive detection (Fig. 5A,

**Figure 5.**
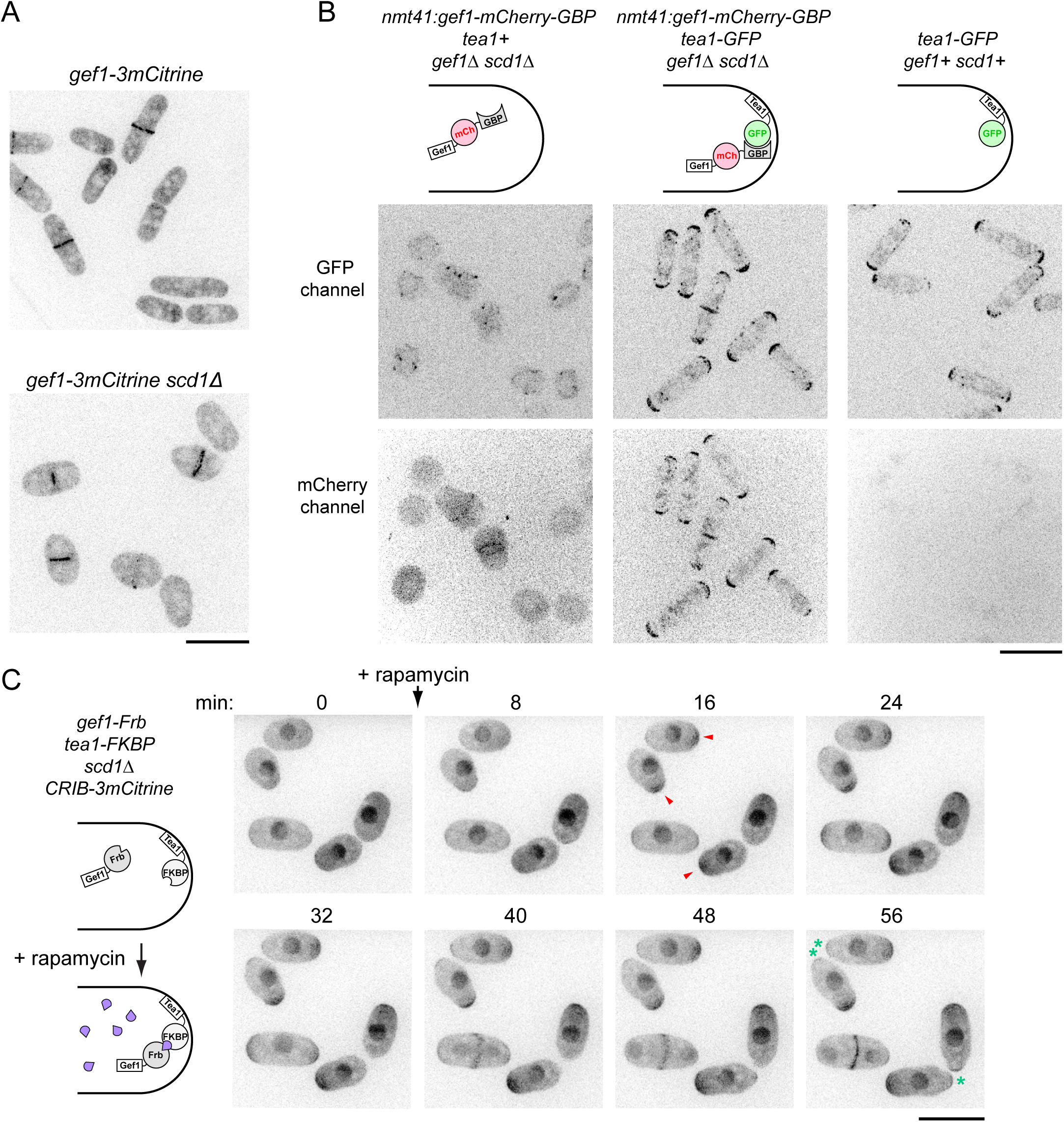
Gef1 is normally cytosolic and is active during interphase, as targeting Gef1 to cell tips in *scd1Δ* cells restores wild-type morphology and Cdc42-GTP enrichment at tips. **(A)** Localization of Gef1-3mCitrine in wild-type and *scd1Δ* cells. In both cases, Gef1 is not at cell tips but is present at the septum. **(B)** Ectopic targeting of Gef1-mCherry-GBP to cell tips by coexpression of Tea1-GFP, in *scd1Δ* background. In cells lacking tagged Tea1 (left-hand panels), Gef1-mCherry-GBP remains cytosolic, and cells are round. In *tea1-GFP* cells (middle panels), Gef1-mCherry-GBP is at cell tips, and cells become strongly polarized. Right-hand panels show absence of bleed-through from GFP channel to mCherry channel. **(C)** Time-resolved targeting of Gef1-Frb to cell tips by rapamycin-induced dimerization in *gef1-Frb tea1-FKBP scd1Δ CRIB-3mCitrine* cells. Rapamycin was added just after the 0 min timepoint. Red arrowheads indicate examples of CRIB-3mCitrine appearance at cell tips after rapamycin addition. Note that after rapamycin addition, cells grow polarized, and eventually CRIB-3mCitrine also appears at opposite cell tips (green asterisks, 56 min), indicating transition from monopolar to bipolar growth. Scale Bars, 10 μm. See also Figure 5--figure supplement 1 and Videos 4 and 5.

Fig. 5—figure supplement 1). We also failed to observe specific localization of Gef1 in interphase *scd1Δ* cells (Fig. 5A,B). Because we recently identified the Sty1 p38 stress-activated MAP kinase pathway as a major regulator of the Cdc42 cell polarity module (Mutavchiev et al., 2016), we hypothesized that cell-tip localization of Gef1 might be associated with cell stress. We therefore treated cells expressing Gef1-3YFP with thiabendazole (TBZ). Like MBC, TBZ depolymerizes MTs; however, unlike MBC, TBZ also leads to a stress that depolarizes the actin cytoskeleton for 60-90 min, via a non-MT-related mechanism (Sawin and Nurse, 1998; Sawin and Snaith, 2004). Interestingly, after treatment with TBZ, Gef1-3YFP transiently localized to cell tips before becoming associated with mobile patches on the plasma membrane on cell sides (Fig. 5—figure supplement 1; Video 4). We conclude that Gef1 is cytosolic in unperturbed interphase cells, and we speculate that mild unintended stresses during some imaging protocols may cause cytosolic Gef1 to associate with the plasma membrane at cell tips (see Discussion).

### Targeting to cell tips converts Gef1 from a global to a local Cdc42 GEF

Together with our finding that *gef1Δ* and *gef1-EANA* mutations restore polarized growth to *tea1Δ scd1*^*low*^ cells, the observation that Gef1 is normally cytosolic suggested that the isotropic growth seen in *scd1* mutant cells in the presence of MBC or in *tea1Δ* or *tea4Δ* backgrounds is due to Gef1 acting on membrane-associated Cdc42 from a cytosolic pool, as a “global” Cdc42 GEF. To support this view, we asked whether artificial targeting of Gef1 to cell tips—that is, changing a “global” Cdc42 GEF into a “local” GEF--would convert it from an activator of isotropic growth into an activator of polarized growth.

In one set of experiments, we used GFP and GFP-binding protein (GBP; (Rothbauer et al., 2008)) to heterodimerize Gef1 with Tea1 (Fig. 5B). Fusion of Gef1-mCherry to GBP rescued the synthetic lethality of *gef1Δ scd1Δ* cells, indicating that Gef1-mCherry-GBP is functional. In *gef1Δ scd1Δ* cells expressing untagged Tea1, Gef1-mCherry-GBP was cytosolic during interphase, and cells displayed the round morphology characteristic of *scd1Δ* mutants. By contrast, in *gef1Δ scd1Δ* cells expressing Tea1-GFP, which is normally localized to cell tips (Behrens and Nurse, 2002), Gef1-mCherry-GBP became localized to cell tips, and cells displayed a normal, wild-type morphology. This demonstrates that targeting Gef1 to cell tips is sufficient to promote highly robust polarized growth in *scd1Δ* cells. In addition, the fact that upon dimerization Gef1 becomes localized to sites of Tea1 localization, rather than vice-versa, supports our finding that Gef1 normally has no specific localization within the cytoplasm.

In a second set of experiments, we used rapamycin-induced dimerization (Chen et al., 1995; Ding et al., 2014; Haruki et al., 2008) to target Gef1 to cell tips (Fig. 5C). Because this method does not require GFP-tagging of Gef1 or its dimerization partner, it allowed us to image CRIB-3mCitrine as a reporter of the Cdc42 polarity module. We first tested dimerization efficiency by replacing endogenous Gef1 and Tea1 with Gef1-Frb-GFP and Tea1-FKBP fusion proteins, respectively. Upon rapamycin addition, Gef1-Frb-GFP rapidly associated with cell tips (Video 5). We then introduced both Gef1-Frb (i.e. without GFP) and Tea1-FKBP fusion proteins into a *scd1Δ CRIB-3mCitrine* background. Before rapamycin addition, interphase cells showed nearly undetectable levels of CRIB-3mCitrine at cell tips. However, upon addition of rapamycin, CRIB-3mCitrine was quickly recruited to cell tips, and morphology and polarized growth became similar to wild-type cells (Fig. 5C).

Taken together with the results above, these results suggest that during interphase, Gef1 is normally localized to the cytosol, where it is active as a global Cdc42 GEF.

### Pom1 kinase activity is required for polarized growth of scd1Δ cells

To understand how MTs, Tea1, and Tea4 may counteract Gef1 to allow polarized growth in *scd1* mutants, we investigated the polarity protein kinase Pom1 (Bahler and Pringle, 1998). Pom1 is localized to the plasma membrane and enriched at cell tips, and this depends both on Tea1 and Tea4 and on Pom1 kinase activity (Hachet et al., 2011). We introduced the analog-sensitive allele *pom1-as1-tdTomato* (Hachet et al., 2011), or control *pom1-tdTomato,* into *scd1Δ GFP-Bgs4 cdc2-asM17* cells and used 3-BrB-PP1 to simultaneously inhibit analog-sensitive Pom1 and Cdc2 (Fig. 6A,B; Video 6). In control *poml-tdTomato* cells, GFP-Bgs4 and Pom1-tdTomato localized to cell tips both before and after 3-BrB-PP1 addition, and cells grew in a polarized manner. In *pom1-as1-tdTomato* cells, GFP-Bgs4 and Pom1-as1-tdTomato localized to cell tips before 3-BrB-PP1 addition; however, after 3-BrB-PP1 addition, both proteins became depolarized, and cells grew isotropically. This demonstrates that Pom1 kinase activity is required for polarized growth of *scd1Δ* cells.

**Figure 6.**
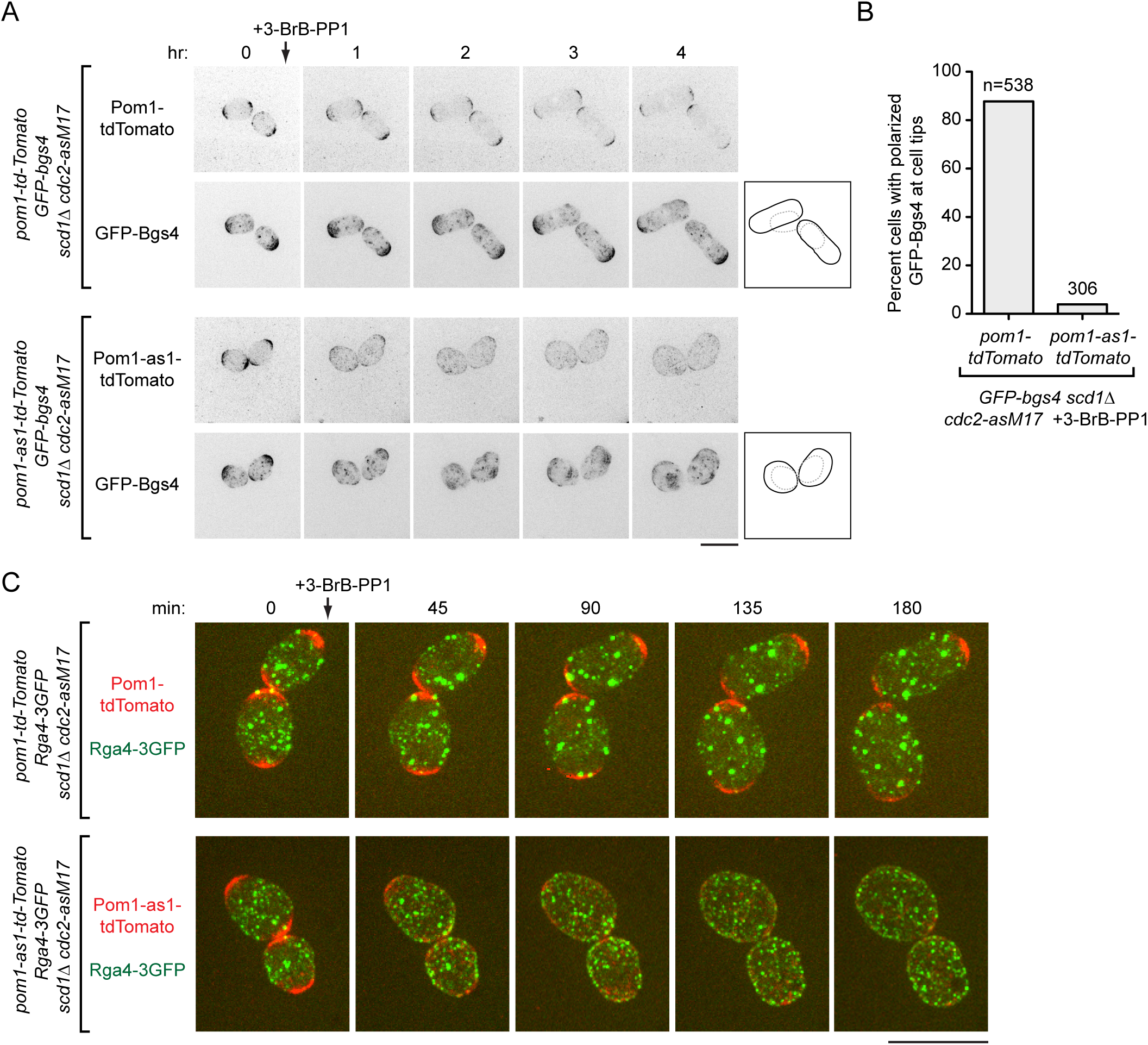
Inhibition of Pom1 kinase activity in *scd1Δ* cells leads to isotropic growth and randomized localization of Cdc42 GAP Rga4. **(A)** Time courses from movies showing cell morphology and distribution of Pom1-tdTomato and GFP-Bgs4, or Pom1-as1-tdTomato and GFP-Bgs4, in *scd1Δ cdc2-asM17* backgrounds after 3-BrB-PP1 treatment. 3-BrB-PP1 inhibits activity of both Cdc2-asM17 and Pom1-as1-tdTomato and was added just after the 0 hr time-point. Diagrams show cell outlines at beginning and end of movies; outlines were aligned slightly because of limited cell movement. Note that 3-BrB-PP1 treatment depolarizes Pom1-as1-tdTomato, and this leads to isotropic growth. **(B)** Percent cells containing polarized GFP-Bgs4 at cell tips, from movies of the type shown in (A). Differences were highly significant (p<0.0001; Fisher’s exact test). **(C)** Time courses from movies showing cell morphology and distribution of Pom1-tdTomato or Pom1-as1-tdTomato, and Rga4-3GFP, in *scd1Δ cdc2-asM17* backgrounds after 3-BrB-PP1 treatment. Scale bars, 10 μm. See also Videos 6 and 7.

To determine whether isotropic growth after Pom1 inhibition depends on Gef1, we introduced either a *pom1Δ* single mutation or *pom1Δ gef1Δ* double mutation into *scd1^low^ mCherry-Bgs4 cdc2-asM17* cells. After 3-BrB-PP1 addition, *pom1A scd1^low^ mCherry-Bgs4 cdc2-asM17* grew isotropically, while *pom1Δ gef1Δ scd1^low^ mCherry-Bgs4 cdc2-asM17* grew in a polarized manner (Fig. 6—figure supplement 1; Video 7). These results suggest that Pom1, like Tea1 and Tea4, contributes to polarized growth of *scd1* mutant cells by counteracting Gef1.

One role of Pom1 is to regulate localization of the Cdc42 GTPase activating protein (GAP) Rga4 (Das et al., 2007; Tatebe et al., 2008). In wild-type cells, Rga4 is localized to the plasma membrane and enriched on cell sides but excluded from cell tips. By contrast, in *pom1Δ* and *pom1* kinase-inactive mutants, Rga4 is no longer excluded from non-growing cell tips (Tatebe et al., 2008). We therefore examined Rga4-3GFP localization in *pom1-as1-tdTomato* and *pom1-tdTomato* cells in *scd1Δ cdc2-asM17* backgrounds after 3-BrB-PP1 addition (Fig. 6C). In control *pom1-tdTomato scd1Δ cdc2-asM17* cells, Rga4-3GFP remained excluded from cell tips. By contrast, in *pom1-as1-tdTomato scd1Δ GFP-Bgs4 cdc2-asM17* cells, Rga4-3GFP quickly became uniformly distributed over the cell surface, coincident with the redistribution of Pom1-as1-tdTomato and the onset of isotropic growth.

### Cdc42 GAP Rga4 counteracts Gef1-dependent isotropic growth

In principle, the uniform surface distribution of Rga4-3GFP after Pom1 inhibition could be either a consequence or a cause of isotropic growth. To distinguish between these possibilities, we investigated how Rga4 contributes to polarized growth when *scd1* function is compromised, and how this is affected by Gef1.

We first analyzed *rga4Δ scd1Δ* double mutants. Previous single-time-point images indicated that *rga4Δ scd1Δ* double mutants are especially wide (Kelly and Nurse, 2011) and, after hydroxyurea arrest, nearly round (Revilla-Guarinos et al., 2016). We introduced *rga4Δ* into *scd1Δ cdc2-asM17 mCherry-Bgs4* cells and imaged cell growth over several hours after 3BrB-PP1 addition. Strikingly, *rga4Δ scd1Δ cdc2-asM17* cells grew completely isotropically, with transient, mobile patches of mCherry-Bgs4 on the cell surface (Fig 7A; Video 8).

**Figure 7.**
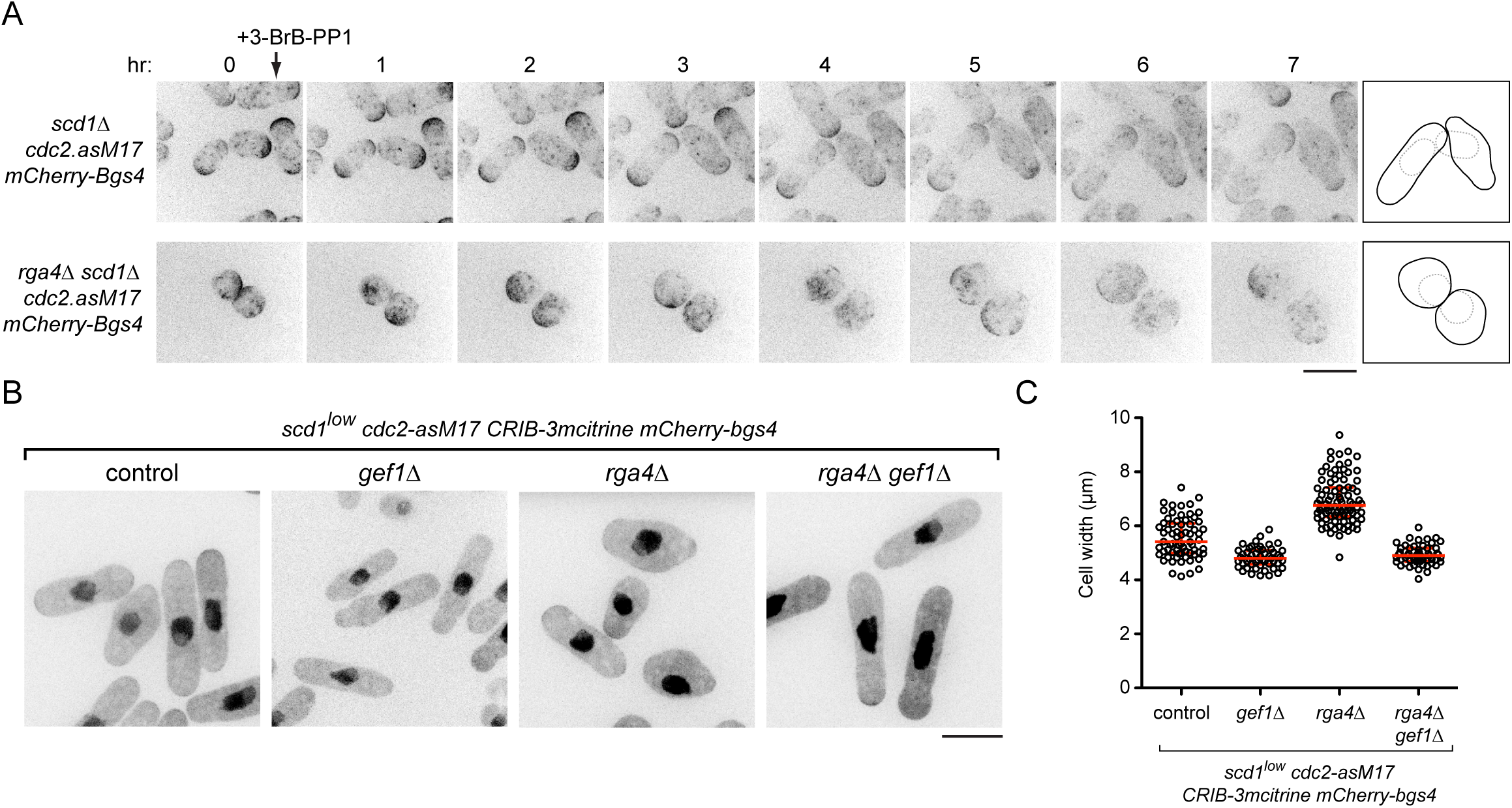
Deletion of *rga4* leads to isotropic growth in *scd1Δ* cells and to Gef1-dependent defects in polarized growth in *scd1*^*low*^ cells. **(A)** Time courses from movies showing mCherry-Bgs4 distribution and cell morphology in *scd1Δ cdc2-asM17 mCherry-bgs4* cells and *rga4Δ scd1Δ cdc2-asM17 mCherry-bgs4* cells after 3-BrB-PP1 treatment. 3-BrB-PP1 was added just after the 0 hr time-point. Diagrams show cell outlines at beginning and end of movies. Note that *rga4Δ scd1Δ* cells grow isotropically; due to mobile nature of mCherry-Bgs4 signal, cell outlines are more clear in movies (see Video 8). **(B)** Morphology of cells with indicated genotypes, in *scd1^low^ cdc2-asM17 CRIB-3mCitrine mCherry-bgs4* background. CRIB-3mCitrine signal is shown here simply to indicate cell outlines. Note that *1Δ* rescues polarity defects associated with *rga4Δ.* **(C)** Quantification of cell width in genotypes shown in B. Median and interquartile ranges are shown. All differences in pairwise comparisons were highly significant (p <0.00001; Student’s t-test, with Bonferroni correction), with the exception of *1Δ* vs. *rga4Δ gef1 A,* which showed no significant difference (p=0.20). For each strain, 51-88 cells were scored. Scale bars, 10 μm. See also Figure 7--figure supplement 1 and Video 8.

In parallel, we introduced *rga4Δ* into *scd1*^*low*^ and *gef1Δ scd1*^*low*^ cells (Fig. 7B,C). The *rga4Δ scd1*^*low*^ cells were compromised in polarized growth, becoming wider and rounder than control *scd1*^*low*^ cells, although this was not as extreme as in *rga4Δ scd1Δ* or *tea1Δ scd1*^*low*^ cells (see Discussion). Importantly, however, these polarity defects were almost completely rescued in *rga4Δ gef1Δ scd1*^*low*^ cells, indicating that the defects associated with *rga4Δ* in *scd1* mutants are indeed mediated through Gef1.

The rescue of *rga4Δ* wide/round morphological defects by *gef1Δ* in a *scd1*^*low*^background appeared to conflict with a previous report that *rga4Δ gef1Δ* double mutants were wider than either *rga4Δ* or *gef1Δ* single mutants (Kelly and Nurse, 2011). We therefore reinvestigated cell dimensions of *rga4Δ* and *gef1Δ* single and double mutants in a wild-type *(scd1+)* background (Fig. 7—figure supplement 1). Consistent with an earlier characterization of *rga4* (Das et al., 2007), we found that *rga4Δ* cells were both shorter and wider at septation compared to wild-type cells. However, we further found that deletion of *gef1* restored *rga4Δ* cells to normal width at septation, and near-normal length. Our results in a wild-type *(scd1+)* background thus contradict previous work (Kelly and Nurse, 2011) and suggest that increased width of *rga4Δ (scd1+)* cells is a consequence of global Gef1 activity competing, albeit with limited success, against relatively strong local Scd1 activity.

## DISCUSSION

### Local and global Cdc42 GEFs

Our results suggest a conceptual model for Cdc42-and MT-mediated cell polarity regulation in fission yeast (Fig. 8) that represents a fundamental departure from previous models (Chang and Martin, 2009; Hachet et al., 2012; Rincon et al., 2014; Sawin and Snaith, 2004)}(Chiou et al., 2017; Kokkoris et al., 2014; Martin and Arkowitz, 2014). While details of the model are presented in Fig. 8, we mention a few key points here.

**Figure 8.**
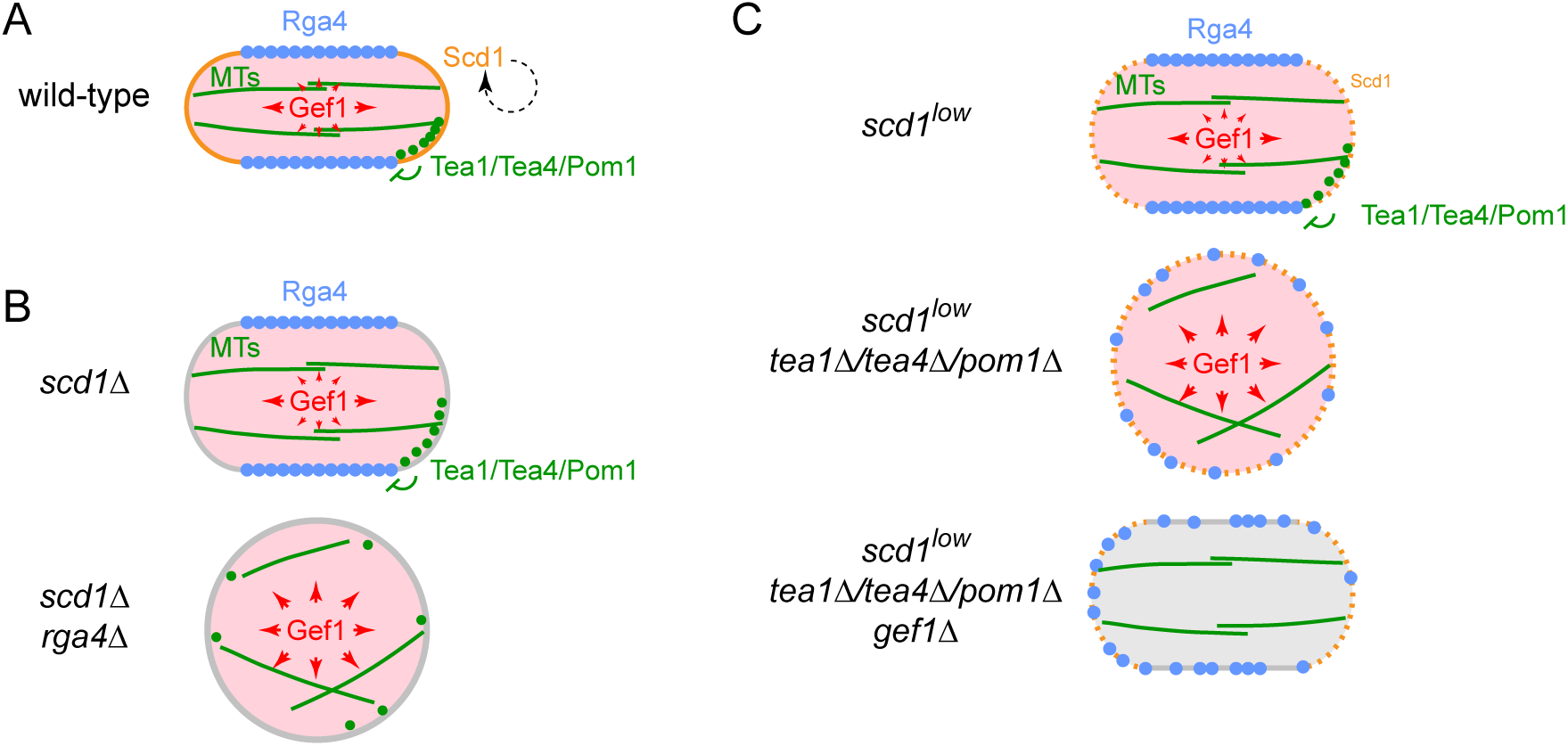
Simplified schematic model for polarized growth via microtubule-dependent coordination of local and global Cdc42 GEF activities. **(A)** In wild-type cells, five main features of the model lead to normal polarized growth: 1) Scd1 (orange) is a plasma membrane-associated “local” Cdc42 GEF at cell tips and maintains a focused polarity zone via positive feedback. 2) Gef1 (pink) is a cytosolic, "global” Gdc42 GEF. 3) Microtubules (MTs; green) target the Tea1/Tea4/Pom1 axis (green) to cell tips. 4) This restricts Cdc42 GAP Rga4 (blue) to the plasma membrane in the cell middle. 5) Rga4 on the membrane locally counters cytosolic Gef1 activity, preventing net GEF activity in the cell middle (represented by large arrows pointing toward cell tips but small arrows pointing elsewhere). **(B)** The model as applied to *scd1Δ* and *scd1Δ rga4Δ* cells. In *scd1Δ* cells, there is no strong focused polarity zone, but Rga4 can still locally counter global Gef1 activity, leading to greater “net” Gef1 activity in the region of cell tips, as in wild-type cells. Therefore, cells are polarized but significantly wider than wild-type. In *scd1Δ rga4Δ cells,* absence of Rga4 means that Gef1 is not locally countered anywhere and is therefore active isotropically. Distribution of MTs and Tea1/Tea4/Pom1 will also ultimately be abnormal, due to round cell shape. **(C)** The model as applied to *scd1^low^, scd1^low^ tea1Δ/tea4Δ/pom1Δ,* and *scd1^low^ tea1ΔJtea4Δpom1Δ gef1Δ* cells. In *scd1^low^cells,* only a very limited amount of the “local” Cdc42 GEF (i.e. Scd1) is present at cell tips, and thus the polarity zone is not focused as in wild-type. However, “net” Gef1 activity remains greater in region of cell tips, and thus Gef1 cooperates with Scd1. In *scd1^low^ tea1Δ/tea4Δ/pom1Δ cells,* Rga4 is no longer spatially restricted. ΔNet” Gef1 activity is therefore isotropic, competing with Scd1 and overwhelming the contribution that low Scd1 levels would otherwise make to polarized growth. In *scd1^low^ tea1Δ/tea4Δ/pom1Δ gef1Δ* cells, competition from Gef1 is alleviated, allowing the limited amount of Scd1 to support polarized growth.

We have shown that Gef1 is a cytosolic, “global” Cdc42 GEF, unlike Scd1, which is a cell tip-localized, “local” Cdc42 GEF (Hirota et al., 2003) (Kelly and Nurse, 2011). Moreover, the functional outputs of these two GEFs are controlled by distinct mechanisms, working in parallel. Promotion of polarized growth by Scd1 is a direct consequence of its localization at cell tips, which is thought to depend on a positive feedback mechanism similar to that in budding yeast (Chiou et al., 2017; Endo et al., 2003; Kelly and Nurse, 2011; Woods and Lew, 2017). By contrast, the spatially uniform cytosolic distribution of Gef1 during interphase would allow it, in principle, to activate Cdc42 anywhere on the plasma membrane. However, this is normally opposed by MTs and the Tea1/Tea4/Pom1 axis, which work to exclude membrane-associated Cdc42 GAP Rga4 from cell tips (Tatebe et al., 2008) and thus “channel” the net Gef1 activity towards cell tips. The importance of restricting net Gef1 activity to cell tips is underscored by our finding that artificial targeting of Gef1 to cell tips in *scd1Δ* cells restores wild-type morphology and CRIB localization at tips.

Previous work by us and others (Bahler and Pringle, 1998; Mata and Nurse, 1997) (Chang and Martin, 2009; Martin et al., 2005; Sawin and Snaith, 2004; Tatebe et al., 2005) suggested that MTs and the Tea1/Tea4/Pom1 axis are important for determining sites of cell polarity establishment, but not for polarity establishment *per se,* or for polarity maintenance. However, the work presented here indicates that this previous view is incomplete: when *scd1* function is compromised, MTs and the Tea1/Tea4/Pom1 axis become essential for polarity maintenance. In addition, our results from *gef1Δ* and loss-of-function *gef1-EANA* cells suggest that the primary role of the Tea1/Tea4/Pom1 axis in polarity maintenance is to counteract, via Rga4, any spatially inappropriate Gef1 activity at cell sides. In mammalian cells, there are similar examples of MTs regulating RhoGEF or RhoGAP distribution or activity, either directly or indirectly, in cell migration, cytokinesis, and tissue organization (Birkenfeld et al., 2008; Etienne-Manneville, 2013; Meiri et al., 2012; Ratheesh et al., 2012; Siegrist and Doe, 2007; Yuce et al., 2005). Although not addressed in the current work, we note that MTs and Tea1/Tea4/Pom1 axis are also important for New-End Take-Off (NETO), the transition from monopolar to bipolar growth (Bahler and Pringle, 1998; Martin et al., 2005; Mata and Nurse, 1997; Mitchison and Nurse, 1985; Nunez et al., 2016).

Our results further suggest that MTs provide the means for coordinating Gef1 function with Scd1 function. Normally, alignment of MTs along the long axis of the cell leads to positioning of MT-dependent landmarks at cell tips (Minc et al., 2009; Terenna et al., 2008) and therefore, ultimately, to enrichment of Rga4 at cell sides. As a result, when MTs and landmarks are present, the Scd1 and Gef1 systems cooperate to promote polarized growth at the same sites, i.e. the cell tips. By contrast, when MTs and/or landmarks are absent, the Scd1 (local) and Gef1 (global) systems can end up competing with each other, with Gef1 promoting isotropic rather than polarized growth (e.g. in *scd1^low^ tea1Δ).*

Our model also provides different mechanistic interpretations of previous results. For example, *scd1Δ* and *rga4Δ* mutations were previously described as having additive effects on cell width, because the *scd1Δ rga4Δ* double mutant was found to be wider than either single mutant (Kelly and Nurse, 2011). However, our work demonstrates that the difference between the single mutants and double mutant is in fact qualitative rather than quantitative. While the two single mutants are both polarized, the double mutant displays a complete loss of polarity and grows isotropically.

### Polarized and isotropic growth in cells with impaired Scd1 function

To analyze polarized growth in *scd1Δ and scd1*^*low*^ cells, we extended the duration of interphase by inhibiting analog-sensitive Cdc2 (Aoi et al., 2014), and we imaged fluorescent-tagged beta-glucan synthase Bgs4, whose localization normally correlates precisely with polarized growth (Cortes et al., 2005). An extended interphase allowed us to unambiguously distinguish polarized vs. isotropic growth in *scd1* mutants, which, because of their short/wide shape, do not normally elongate very much during a single cell cycle. In addition, imaging during extended interphase can circumvent problems that may arise when strains have abnormal phenotypes associated with cytokinesis (e.g. *pom1Δ;* (Bahler and Pringle, 1998) see Methods). Interestingly, during isotropic growth, Bgs4 appeared as transient and mobile patches on the plasma membrane instead of being distributed homogeneously. The transient nature of these patches will be interesting to investigate in the future.

The use of analog-sensitive Cdc2 is not expected to affect the interpretation of our results. In fission yeast, polarized growth continues when Cdc2 kinase is inactivated by either temperature-or analog-sensitive mutations (Dischinger et al., 2008; Nurse et al., 1976). In the absence of inhibition, *cdc2-asM17* retains essentially all of the functionality of wild-type *cdc2+* (Aoi et al., 2014), unlike an earlier *cdc2-as* allele (Dischinger et al., 2008). Moreover, to inhibit Cdc2-asM17, we used the minimum concentration of analog required to prevent mitotic entry (see Methods). Under these conditions (i.e. in the absence of any other perturbations), both wild-type and *scd1Δ* cells show robust polarized growth--without, for example, any noticeable increase in cell width.

While our initial experiments involved *scd1Δ* cells, many subsequent experiments involved *scd1*^*low*^ cells. This was essential for deciphering the relationship between Scd1,

Gef1 and the Tea1/Tea4/Pom1 axis, because *scd1Δ* is synthetically lethal with *tea1Δ* and *gef1Δ,* while *scd1*^*low*^ is not. At the same time, this difference in synthetic lethality highlights the fact that because *scd1*^*low*^ cells retain some Scd1 function, they are not equivalent to *scd1Δ* cells. In particular, *scd1Δ rga4Δ* cells show strongly isotropic growth, while *scd1^low^ rga4Δ* cells have less severe polarity defects (which are nevertheless rescued by *gef1Δ).* The simplest explanation for this is that in *scd1^low^ rga4Δ* cells, the polarity system set up by low levels of Scd1 can partially compete against the Gef1-dependent drive towards isotropic growth. How low levels of Scd1 achieve this at a mechanistic level remains to be explored.

In this context, it is also interesting to compare polarity phenotypes of *scd1^low^ rga4Δ* with *scd1^low^ tea1Δ,* because *scd1^low^ tea1Δ* cells grow completely isotropically (as do *scd1^low^ tea4Δ,* and *scd1^low^ pom1Δ),* even though they express enough Scd1 to maintain viability in a *tea1Δ* genetic background. We can imagine two non-exclusive explanations for this difference in phenotype. First, in addition to regulating Rga4, the Tea1/Tea4/Pom1 axis could have a separate role in either bolstering *scd1*^*low*^ function or countering *gef1* function. Second, the different phenotypes could be due to the presence vs. the absence of Rga4. That is, in *scd1^low^ tea1Δ* cells, the GAP activity of Rga4 will be distributed essentially evenly over the entire plasma membrane, including at “prospective tip” regions, thereby counteracting the weak polarizing activity of Scd1^low^; by contrast, in *scd1^low^ rga4Δ* cells, there is no Rga4 GAP activity anywhere, and therefore low levels of Scd1 may have a greater net effect on cell polarity.

### Gef1 localization and regulation in unstressed cells

One important difference between our work and existing literature (Das et al., 2015; Das et al., 2009; Kokkoris et al., 2014; Vjestica et al., 2013) is that we find that in unperturbed interphase cells, Gef1 is not present at cell tips but is rather a cytosolic, global Cdc42 GEF, both in wild-type and in *scd1Δ* cells. It is unclear why several previous reports observe Gef1 at interphase cell tips. Our own results lead us to speculate that this may be due to unintended mild cell stress, possibly because of how cells are prepared for imaging, or because of phototoxicity during imaging (Laissue et al., 2017). In our experiments, cells are imaged under conditions that are essentially identical to those of cells growing exponentially in flasks, apart from shaking. This minimizes stress ((Mutavchiev et al., 2016); see Methods) and allows imaging of polarized growth under the microscope for several hours.

Previous work has suggested that Gef1 is negatively regulated by phosphorylation via the NDR kinase Orb6 (Das et al., 2015; Das et al., 2009); specifically, Orb6 is thought to prevent Gef1 from localizing to the plasma membrane on cell sides. Our results are not inconsistent with this view. However, because we find that Gef1 can be active as a cell-polarity GEF from the cytosol, we would argue that the regulation of Gef1 membrane localization (specifically, to cell sides) is separable from the regulation of Gef1’s GEF activity *per se.* It is possible that localization of Gef1 to the plasma membrane on cell sides might further potentiate its net biological activity relative to any countering GAP activity from Rga4. These will be interesting questions to address in the future.

Regulated localization of Cdc42 GEFs to the plasma membrane may also be relevant to mammalian cells. Gef1 is unusual among RhoGEFs in that while it contains a catalytic DH domain, it lacks a pleckstrin homology (PH) domain, which is present in nearly all DH-family RhoGEFs and is important for association with membrane lipids (Cook et al., 2014; Rossman et al., 2005). The mammalian Cdc42 GEF Tuba also lacks a PH domain and instead contains a BAR domain (Salazar et al., 2003); Gef1 has also been proposed to contain a BAR domain, although this has not been confirmed experimentally (Das et al., 2015). Interestingly, in MCDK epithelial cells, Tuba is localized to the cytoplasm when cells are grown in a monolayer but is concentrated subapically when cells are grown to form cysts (Qin et al., 2010). Thus, like Gef1, the localization of Tuba may be subject to regulation, during development/differentiation.

### Concluding remarks

What might be the purpose of regulating cell polarity by both local and global Cdc42 GEFs? While here we can only speculate, we note that *gef1Δ* cells have a mild defect/delay in NETO (Coll et al., 2003; Das et al., 2012). Computational modeling suggests that Gef1’s contribution to total Cdc42 GEF activity may be an important feature in the timing of NETO and in the symmetry of Cdc42 activation at the two cell tips (Das et al., 2012). In light of our results, it would may be of interest to investigate, in a more detailed spatial model, how the particular properties of a local vs. a global GEF may influence the NETO transition. A second possible purpose relates to our observation that although Gef1 is cytosolic in unperturbed cells, it associates with the plasma membrane upon TBZ stress (this work), as well as upon inhibition/inactivation of Orb6 (Das et al., 2009). Thus, Gef1 may have a specific role in regulating cell polarity in response to stress.

## METHODS

### Yeast culture

Standard fission yeast methods were used throughout (Forsburg and Rhind, 2006; Petersen and Russell, 2016). Growth medium was either YE5S rich medium (using Bacto yeast extract; Becton Dickinson) or PMG minimal medium, with glucose added after autoclaving (PMG is equivalent to EMM2 minimal medium, but uses 4 g/L sodium glutamate instead of ammonium chloride as nitrogen source). PMG was used only for experiments involving *scd1*^*low*^ cells (i.e. *nmt81:3HA-scd1* cells), in which case cells were grown first in PMG (i.e. without thiamine) and then in PMG plus 20 μM thiamine for 24 hr prior to use in imaging experiments. In all other experiments (i.e. all experiments not involving *scd1*^*low*^ cells) YE5S was used. Supplements such as adenine, leucine, and uracil were used at 175 mg/L. Solid media used 2% Bacto agar (Becton Dickinson).

We note that when grown on solid PMG medium without thiamine, *scd1^low^ tea1Δ, scd1^low^ tea4Δ, and scd1^low^ pom1Δ* double mutants formed colonies that were noticeably smaller than wild-type cells and *scd1*^*low*^ single mutants. Under these conditions, the double mutants also showed some defects in septum positioning and in completion of cytokinesis.

### Plasmid and yeast strain construction

Mating for genetic crosses (Ekwall and Thon, 2017) was performed on SPA5S plates with supplements at 45 mg/L. Crosses were performed using tetrad dissection or random spore analysis. Tagging and deletion of genes were performed using PCR-based methods (Bahler et al., 1998), with the exception of the strains described below, which involved integration of newly-constructed plasmids. All plasmid constructions (below) were confirmed by sequencing. All strains used in this study are listed in Supplementary File 1.

*adh13:wee1* plasmid/strain construction. The *wee1* open reading frame (ORF) was amplified by PCR from genomic DNA and cloned into the NdeI site of pNATZA13 (kind gift from Y. Watanabe) to form pNATZA13-Wee1 (pKS1448). ApaI-linearised pKS1448 was then integrated at the Z locus (Sakuno et al., 2009) of KS515, and positive clones were screened by microscopy and confirmed by colony PCR.

*gef1-EANA-3mCherry.* TOPO-Gef1-3mCherry:kan plasmid (pKS1632) was constructed using 3-piece Gibson assembly approach (NEB). Briefly, PCR fragments of TOPO vector (pCR2.1), Gef1 ORF (flanked by 180bp upstream of Gef1 ORF), and 3mCherry-Kan fragment (flanked by 180bp downstream of Gef1 ORF) were assembled to generate pKS1632. A PCR fragment of Gef1 (internal fragment corresponding to amino-acid residues 314-508 but containing two point mutations, E318A and N505A) was subsequently introduced into pKS1632 via a 2-piece Gibson assembly approach to generate pKS1699. A Gef1-containing SpeI-XbaI fragment from pKS1699 was then purified and transformed into strain KS7656 to generate strain KS9183. gef1-mCherry-GBP. The *gef1+* ORF was amplified from genomic DNA and introduced into pINTH41.3HA-mCherry-GBP-3PK:natMX6 plasmid (kind gift from I. Hagan) via 2-piece Gibson assembly to generate pKS1488. NotI-linearised pKS1488 was then transformed into strain KS7742 to generate strain KS8152.

### Cell lysate preparation and western blotting

For measuring 3HA-Scd1 levels by Western blotting, cell lysates were prepared using the trichloroacetic acid (TCA) lysis protocol (Grallert and Hagan, 2017). For each sample, approximately 5 × 10^7^ cells (10 mL culture at 5×10^6^ cells/mL) in PMG liquid media (or PMG plus thiamine, if after thiamine addition) in a 25°C shaking water bath were collected by centrifugation, washed with 1 mL stop buffer, transferred to a 1.5 mL screw-cap tube, and flash-frozen in liquid nitrogen. Then, 200 μl of ice-cold 20% TCA and approximately 400 μl of 0.5 mm zirconium/silica beads (BioSpec; 11079105z) were added to the cell pellet. Cell lysis was performed in a Ribolyser bead-beater (Hybaid) using one cycle of 45 sec at speed setting “6”. 400 μl of 5% TCA was then added to the suspension, and the cell lysate was recovered by puncturing the bottom of the screw cap tube and centrifuging into a 1.5 mL microfuge tube (5 min, 13k rpm, 4°C). The TCA supernatant was discarded, and the protein pellet was then solubilized in Laemmli sample buffer and incubated at 100°C for 3 minutes. Solubilized lysates were separated by SDS-PAGE and transferred by western blotting to 0.2 μm nitrocellulose filters (Biorad). Western blots were stained with Ponceau S and scanned prior to blocking. Blocking and antibody incubations used 5% milk in TBS with 0.05% Tween 20. 12CA5 anti-HA primary antibody (Roche; 11583816001, used at 1:1000 dilution) and IRDye 680CW Donkey anti-Mouse secondary antibody (Licor; 926-68022, 1:10000) were used to probe 3HA-Scd1. Blots were imaged using an Odyssey fluorescence imager (Licor) and processed using Image Studio Lite (Licor). Two biological replicates of Western blots were performed.

### Microscopy sample preparation and imaging

All imaging experiments were performed with exponentially growing cells cultured at 25°C. Imaging was performed either in coverslip dishes (MatTek; P35G-0.170-14-C.s) or 4-chamber glass bottom micro-slides (Ibidi; 80427). Imaging dishes/slides were placed on a 25°C heat block, coated with 1 mg/mL soybean lectin (Sigma; L1395), left for 10 min, and washed with appropriate medium to remove excess lectin. Log-phase culture was added to dishes/slides and left to settle for 15 minutes. The dishes/slides were washed extensively with media using aspiration with at least 3 full exchanges of media (approximately 1 mL each). Finally, 500 μL of medium was added to the dish/slide before imaging.

Live-cell fluorescence imaging was performed using a custom spinning-disk confocal microscope unit [Nikon TE2000 microscope base, attached to a modified Yokogawa CSU-10 unit (Visitech) and an iXon+ Du888 EMCCD camera (Andor), 100x/1.45 NA Plan Apo objective (Nikon), Optospin IV filter wheel (Cairn Research), MS-2000 automated stage with CRISP autofocus (ASI), and thermo-regulated chamber maintained at 25°C (OKOlab)]. Metamorph software (Molecular Devices) was used to control the spinning-disc confocal microscope.

3-BrB-PP1 (4-Amino-1-tert-butyl-3-(3-bromobenzyl)pyrazolo[3,4-d]pyrimidine; A602985) was obtained from Toronto Research Chemicals and dissolved in methanol to make a 50 mM stock solution. 3-BrB-PP1 was used at a final concentration of 8 pM; 4 pM was insufficient to completely prevent mitotic entry. Thiamine, methyl-2-benzimidazole carbamate (MBC), and thiabendazole (TBZ) were obtained from Sigma. Thiamine was dissolved in water as 200 mM stock and used at a final concentration of 20 μM. MBC stock solution was 2.5 mg/mL in DMSO and was used at a final concentration of 25 μg/mL (therefore 1% DMSO final concentration). In MBC experiments, DMSO-only control used 1% DMSO final. TBZ stock solution was 30 mg/mL in DMSO and was used at a final concentration of 150 μg/mL (therefore 0.5% DMSO final concentration). Rapamycin was obtained from Fisher Scientific Ltd (Cat No. 10798668). Rapamycin was dissolved in DMSO as a 1 mg/mL stock and used at a final concentration of 2.5 pg/mL All drug additions during imaging were performed by medium exchange using a 1 mL polyethylene transfer pipette (Fisher Scientific, 1346-9118).

Numbers of independent biological replicate experiments are provided for each yeast strain, in each figure, in the yeast strain list in Supplementary File 1. We define an independent biological replicate as growing/culturing a given yeast strain and then using it for a given biochemistry experiment or imaging session. Up to a few hundred cells of the same genotype may be imaged in any given replicate imaging session.

### Analysis of microscopy images

Processing of the acquired raw images was executed using ImageJ (Fiji, NIH).

Unless otherwise stated, all images and videos shown are maximum projections of eleven Z-sections with 0.7 μm step-size. For rigid body registrations, ImageJ StackReg and Linear Stack Alignment with SIFT plugins were used. Image formatting and assembly were performed using Photoshop (Adobe) and Illustrator CS3 (Adobe). Cell outlines were drawn by hand in Illustrator, using images from individual video time-points as templates. In a few cases where cell borders were more difficult to discern (e.g. in late-stage depolarized cells), images from successive time-points were superimposed and then used as a template for drawing. Videos were edited using ImageJ and QuickTime (Apple).

Quantification of the percentage of cells with polarized mCherry-Bgs4 or GFP-Bgs4 signals on cell tips (Fig 2, 3, 4 and 6) was performed manually, based on analysis of videos. Cells with persistent mCherry-Bgs4 or GFP-Bgs4 signals on the cell tips (over a period of four hr) were scored as polarized cells. To avoid confusing depolarized cells with cells that simply had a diminished Bgs4 signal (because of photo-bleaching), quantification was performed using only on the first four hr of videos. Occasional cells that transiently lost the tip signal but then regained it shortly afterwards (i.e. in the same place) were also scored as polarized cells. Graphs were created using Graphpad Prism software. Statistical analysis was carried out using online tools (<u>http://www.graphpad.com/quickcalcs/</u>; <u>http://www.socscistatistics.com</u>).

## ACKNOWLEDGEMENTS

We thank Mohan Balasubramanian, Susan Forsburg, Iain Hagan, Heinrich Leonhardt, Sophie Martin, Snezhana Oliferenko, Juan Carlos Ribas, Masamitsu Sato, Kazuhiro Shiozaki, and Hilary Snaith for strains and/or reagents, Kent Rossman for fission yeast RhoGEF alignment and advice on design of *gef1-EANA,* and members of our laboratories for discussion and encouragement.

This work was supported by the BBRSC [BB/K021699/1] and the Wellcome Trust [094517]. The Wellcome Centre for Cell Biology is supported by core funding from the Wellcome Trust [203149].

## LEGENDS TO VIDEOS

### Video 1. Microtubule depolymerization in *scd1Δ* cells leads to isotropic growth

mCherry-Bgs4 distribution and cell morphology of *scd1Δ cdc2-asM17 mCherry-bgs4* cells. Bgs4 on the plasma membrane indicates sites of growth. Cells were pre-treated with 3-BrB-PP1 60 min prior to start of imaging, to inhibit Cdc2 kinase activity, and then treated with either DMSO or MBC at start of imaging. For DMSO treatment, cell at lower right corresponds to cell shown in Fig. 2A. For MBC treatment, cell at mid-lower center corresponds to cell shown in Fig. 2A. Time interval during acquisition, 10 min; total elapsed time, 420 min; time compression at 15 frames per second playback, 9000X.

### Video 2. When scd1 is expressed at very low levels, tea*1Δ* and tea4Δ cells grow isotropically

mCherry-Bgs4 distribution and cell morphology in control cells, *tea1Δ,* and *tea4Δ* cells, all in a *scd1^low^ cdc2-asM17 mCherry-bgs4* genetic background. Bgs4 on the plasma membrane indicates sites of growth. Cells correspond to those shown in Fig. 3. *scd1* expression was repressed by thiamine addition 24 hr prior to start of imaging. Cdc2 kinase activity was inhibited by 3-BrB-PP1 addition 30 min before imaging. Time interval during acquisition, 10 min; total elapsed time, 420 min; time compression at 15 frames per second playback, 9000X.

### Video 3. Loss of *gef1* function restores polarized growth to *scd1^low^ tea1Δ* cells

mCherry-Bgs4 distribution and cell morphology in *tea1Δ, gef1Δ., tea1Δ gef1Δ* and *tea1Δ gef1-EANA* cells, all in a *scd1^low^ cdc2-asM17 mCherry-bgs4* genetic background. Bgs4 on the plasma membrane indicates sites of growth. Cells correspond to those shown in Fig. 4A. *scd1* expression was repressed by thiamine addition 24 hr prior to start of imaging. Cdc2 kinase activity was inhibited by 3-BrB-PP1 addition 30 min before imaging. A transient loss of Bgs4 from cell tips is seen in some *scd1^low^ gef1* mutant cells (~20%), including some of the examples shown in the movie; the reasons for this are not completely clear. Time interval during acquisition, 10 min; total elapsed time, 420 min; time compression at 15 frames per second playback, 9000X.

### Video 4. Gef1-3YFP is transiently recruited to the cell tips upon TBZ treatment

TBZ was added just after the first time-point, which is paused in the movie. Prior to TBZ addition, Gef1-3YFP in dividing cells is present at the division site and in the cytoplasm, and Gef1-3YFP In interphase cells is uniformly distributed in the cytoplasm, without any visible enrichment at the cell tips. Upon TBZ addition, interphase Gef1-3YFP signal is transiently observed at cell tips and later appears to move along the cell cortex towards the cell middle. Three of the cells in the movie correspond to those shown in Fig. 5—figure supplement 1. Time interval during acquisition, 9 min; total elapsed time, 81 min; time compression at 15 frames per second playback, 8100X.

### Video 5. Gef1-Frb-GFP is recruited to cell tips via interaction with Tea1-2FKBP upon rapamycin addition

Rapamycin was added just after the first time-point, which is paused in the movie. Prior to rapamycin addition, Gef1-Frb-GFP in dividing cells is present at the division site and in the cytoplasm, and Gef1-Frb-GFP in interphase cells is uniformly distributed in the cytoplasm, without any visible enrichment at the cell tips. Upon rapamycin addition, Gef1-Frb-GFP is rapidly recruited to the cell tips in a pattern closely resembling Tea1 localization (see Fig. 5B). Time interval during acquisition, 5 min; total elapsed time, 35 min; time compression at 15 frames per second playback, 4500X.

### Video 6. Inhibition of Pom1 kinase activity in *scd1Δ* cells leads to isotropic growth

Cell morphology and distribution of Pom1-tdTomato and GFP-Bgs4, or Pom1-as1-tdTomato and GFP-Bgs4, in *scd1Δ cdc2-asM17* genetic background after 3-BrB-PP1 treatment. Bgs4 on the plasma membrane indicates sites of growth. Cells correspond to those shown in Fig. 6A. 3-BrB-PP1 inhibits activity of both Cdc2-asM17 and Pom1-as1-tdTomato and was added just after the first time-point. Note that 3-BrB-PP1 treatment depolarizes Pom1-as1-tdTomato, and this leads to isotropic growth. Time interval during acquisition, 20 min; total elapsed time, 240 min; time compression at 15 frames per second playback, 18,000X.

### Video 7. Deletion of *gef1* restores polarized growth to *scd1^low^pomlA* cells

mCherry-Bgs4 distribution and cell morphology of *pom1Δ* and *pomΔ gef1Δ* mutants in *scd1^low^ cdc2-asM17 mCherry-bgs4* genetic background. Bgs4 on the plasma membrane indicates sites of growth. Cells correspond to those shown in Fig. 6—figure supplement 1. *scd1* expression was repressed by thiamine addition 24 hr prior to start of imaging. Cdc2 kinase activity was inhibited by 3-BrB-PP1 addition 30 min before imaging. Time interval during acquisition, 10 min; total elapsed time, 350 min; time compression at 15 frames per second playback, 9000X.

### Video 8. Deletion of *rga4* leads to isotropic growth in *scd1Δ* cells

mCherry-Bgs4 distribution and cell morphology of *scd1Δ* and *scd1Δ rga4Δ* mutants in *cdc2-asM17 mCherry-bgs4* background. Bgs4 on the plasma membrane indicates sites of growth. Cells correspond to those shown in Fig. 7A, with slightly larger fields. Cdc2 kinase activity was inhibited by 3-BrB-PP1 addition 30 min before imaging. Time interval during acquisition, 12 min; total elapsed time, 480 min; time compression at 15 frames per second playback, 10,800X.

## FIGURE SUPPLEMENTS

Figure 1—figure supplement 1. Reporters of the Cdc42 polarity module are not detected at cell tips in polarized *scd1Δ* cells.

Figure 3—figure supplement 1. Thiamine-mediated scd1 repression via *nmt41* and *nmt81* promoters.

Figure 4—figure supplement 1. *gef1-EANA-3mCherry* is a *gef1* loss-of-function mutation.

Figure 5—figure supplement 1. Gef1 is cytosolic except under stress conditions.

Figure 6—figure supplement 1. Deletion of *gef1* restores polarized growth to *scd1^low^ pom1Δ* cells.

Figure 7—figure supplement 1. *gef1Δ* rescues the short/wide phenotype of *rga4Δ* cells.

**Figure 1--figure supplement 1.**
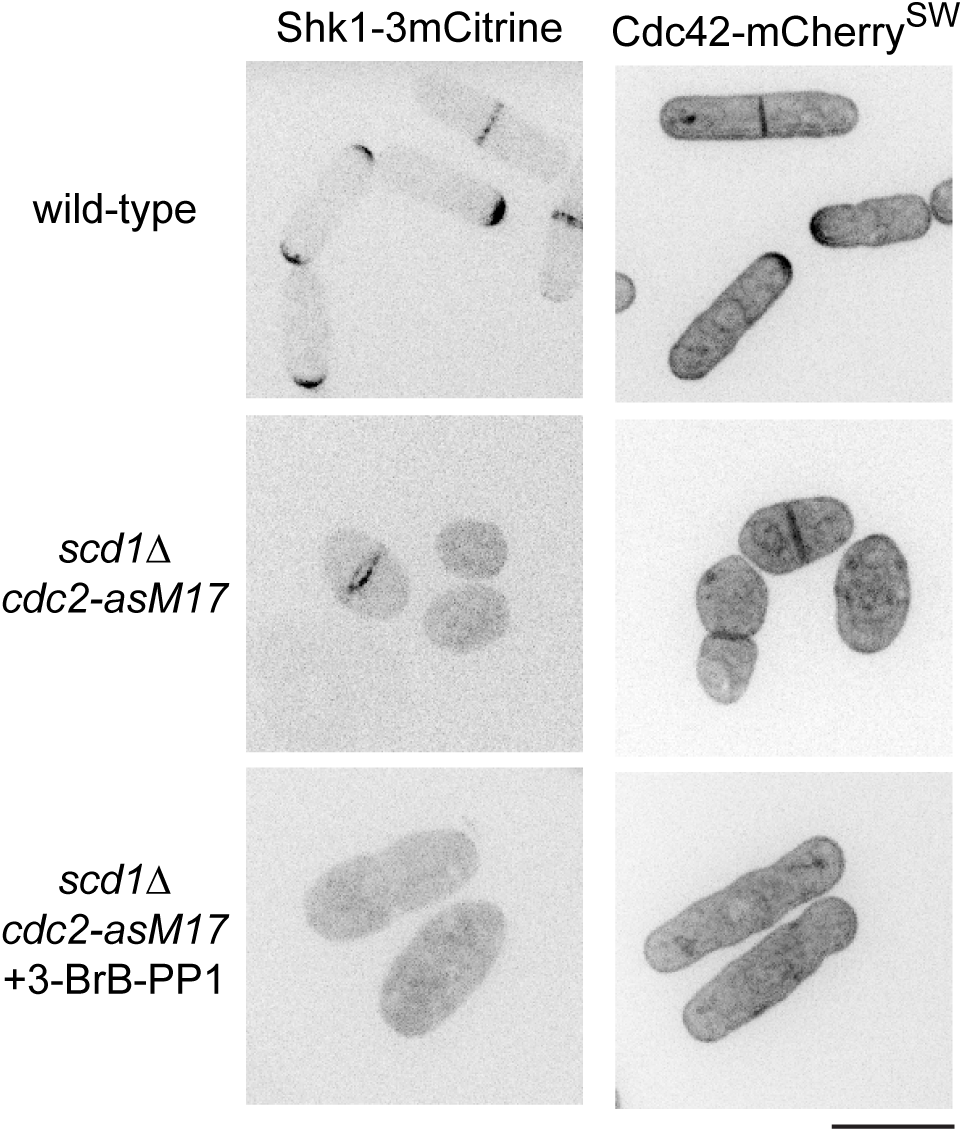
Reporters of the Cdc42 polarity module are not detected at cell tips in polarized. *scd1Δ* **cells.** Localization of additional reporters of the Cdc42 polarity module (i.e. other than the CRIB-3mCitrine reporter shown in Fig. 1), in cells of the indicated genotypes. In polarized *scd1Δ* cells (bottom panels), Shk1-3mCitrine is not detected at cell tips, and Cdc42-mCherry^SW^ is not enriched at cell tips, even though it remains associated with the plasma membrane. Unlike the high-contrast CRIB reporter, these additional reporters do not show any nuclear localization; therefore, their absence from cell tips in *scd1Δ* cells cannot be attributed to an increased nuclear localization. In bottom panels, cells were treated with 3-BrB-PP1 for 5 hr prior to imaging. Scale bar, 10 μm.

**Figure 3--figure supplement 1.**
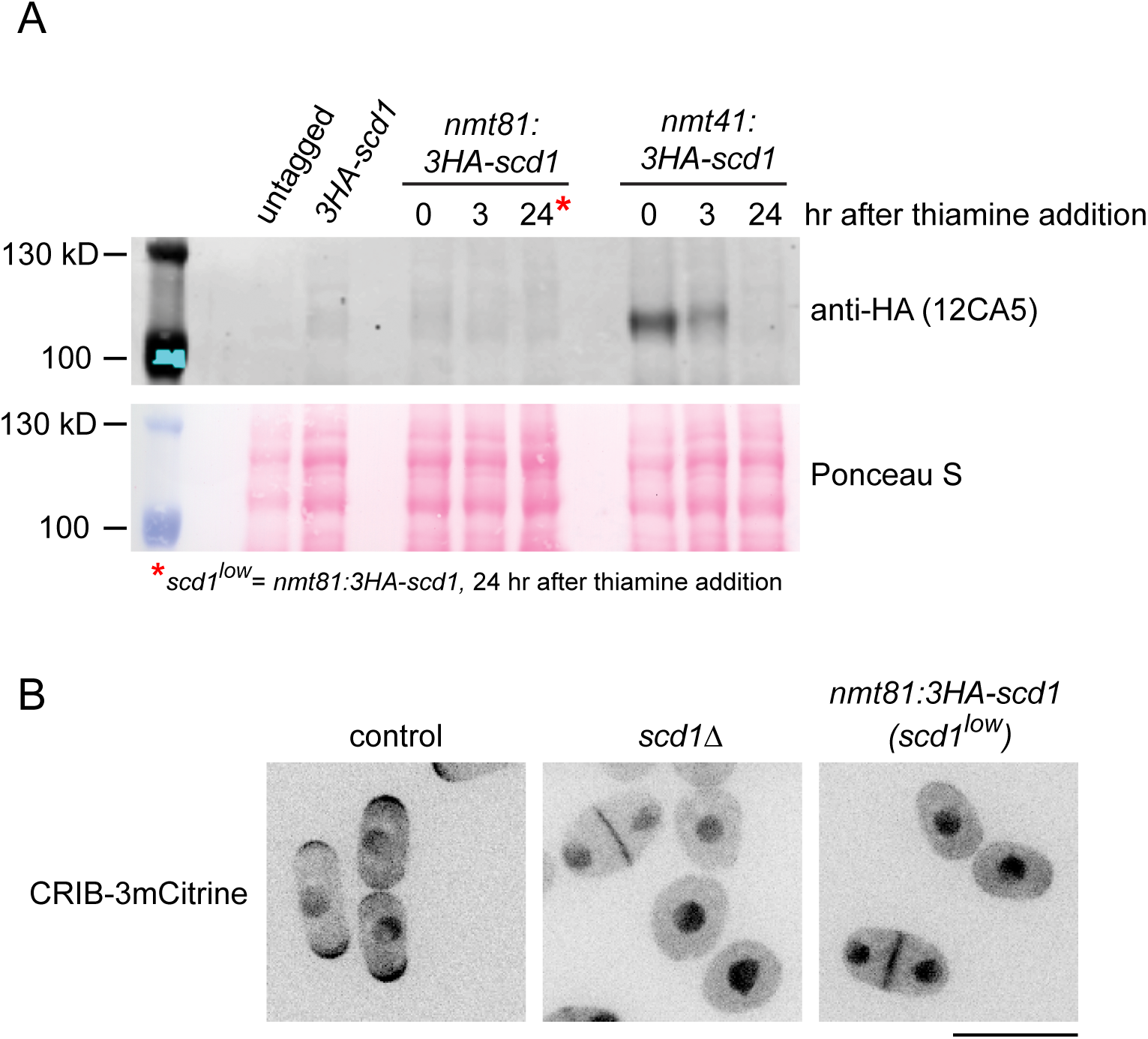
. Thiamine-mediated *scd1* repression via *nmt41* and *nmt81* promoters. **(A)** Anti-HA Western blot of lysates from untagged cells and from cells expressing 3HA-Scd1 from the endogenous scd1 promoter (3HA-scd1), from the weak nmt81 promoter (nmt81:3HA-scd1), and from the medium-strength nmt41 promoter (nmt41:3HA-scd1), before and after repression by thiamine addition. In main text, nmt81:3HA-scd1 cells after 24 hr of repression are referred to as μscd1^*low*^ cells”. Under these conditions, 3HA-Scd1 is barely detectable above background. **(B)** Morphology and CRIB-3mCitrine localization in control, scd*1Δ,* nmt81:3HA-scd1 cells in a cdc2-asM17 genetic background (in this instance, without inhibition of Cdc2-asM17 by 3-BrB-PP1). All cells were grown in PMG liquid media. Expression in nmt81:3HA-scd1 cells (scd1^*low*^ cells) was repressed by 20 μM thiamine for 24 hours prior to imaging. Note that as in scd*1Δ* cells, in nmt81:3HA-scd1 cells (scd1^*low*^ cells), CRIB-3mCitrine is not detected at the tips of interphase cells. Scale bar, 10 pm.

**Figure 4--figure supplement 1.**
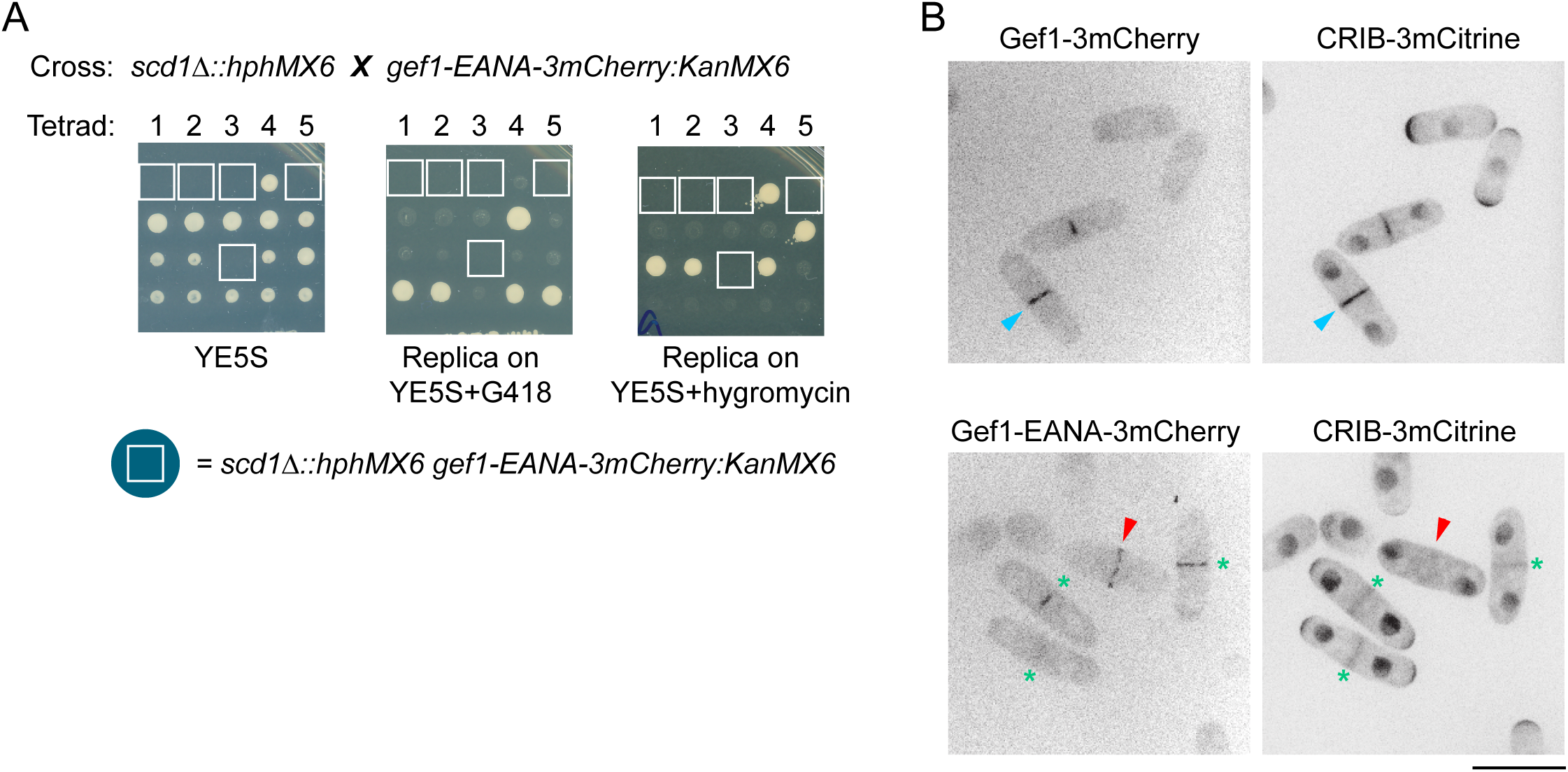
*gef1-EANA-3mCherry* is a *gef1* loss-of-function mutation. **(A)** Tetrad analysis showing synthetic lethality of *gef1-EANA-3mCherry* with *scd1Δ,* similar to the known synthetic lethality of *gef1Δ* with *scd1Δ* (Coll et al., 2003). Spores were germinated on YE5S, and resulting colonies replica-plated to YE5S+G418 and YE5S+hygromycin. Tetrads 1,2 and 5 are tetratypes; tetrad 4 is a parental ditype; tetrad 3 is a non-parental ditype. Boxes indicate inferred position of non-viable *gef1-EANA-3mCherry scd1Δ* double mutants. **(B)** In septating cells, both wild-type Gef1-3mCherry and mutant Gef1-EANA-3mCherry localize to the septation region. However, in early stages of septation, wild-type Gef1-3mCherry recruits Cdc42-GTP, indicated by CRIB-3mCitrine (blue arrowheads) while Gef1-EANA-3mCherry does not recruit CRIB-3mCitrine (red arrowheads). In later stages of septation, CRIB-3mCitrine in the septation region is more diffuse and weak and does not correlate with Gef1-EANA-3mCherry (green asterisks); this later localization is known to be independent of Gef1 (Wei et al., 2016). Scale bar, 10 pm.

**Figure 5--figure supplement 1.**
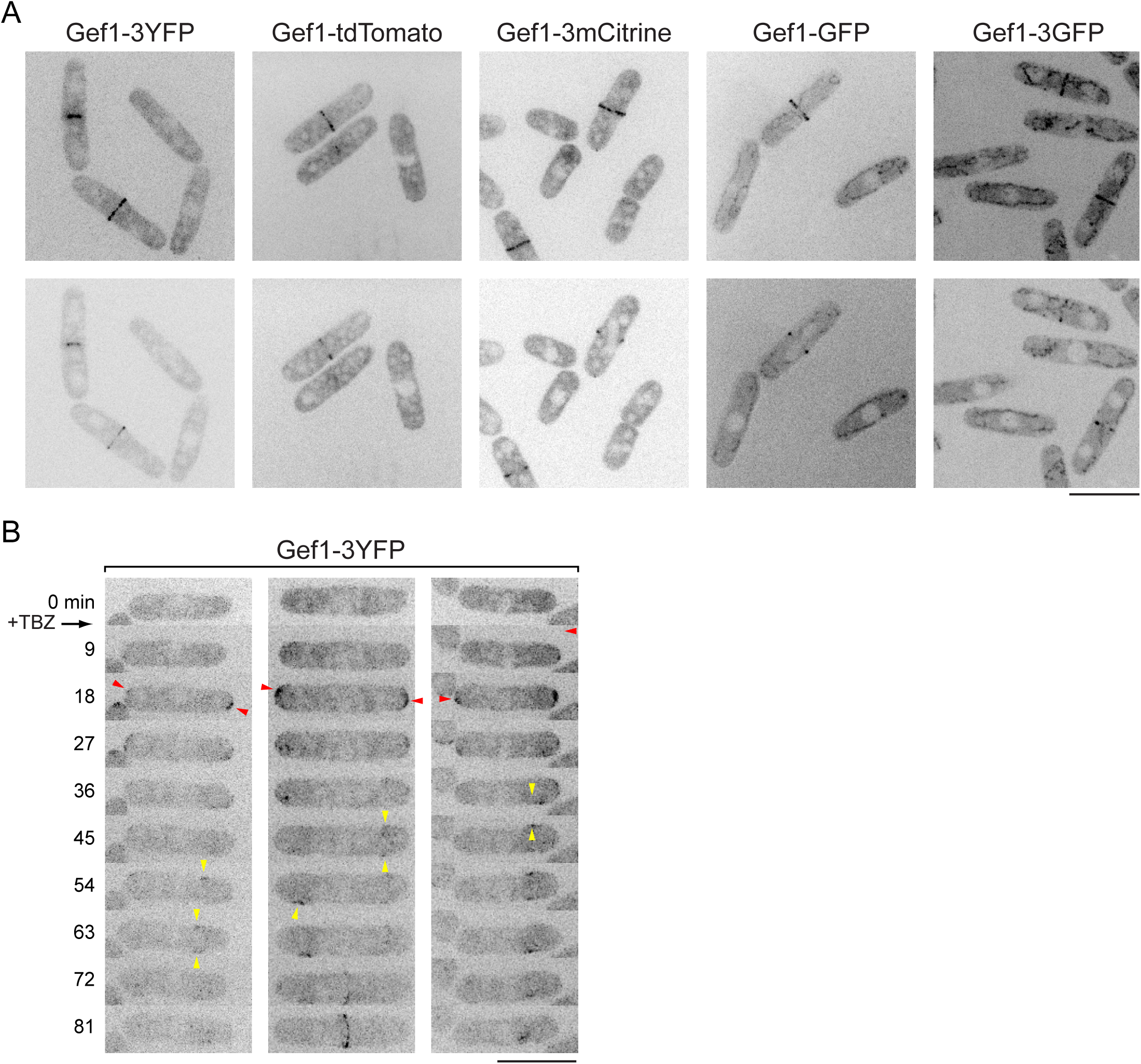
Gef1 is cytosolic except under stress conditions. *(A)* Still images of Gef1 fused to different fluorescent proteins. Top panels show maximum projections, and bottom panels show corresponding central Z-section or two adjacent central Z-sections. During interphase, Gef1 is cytosolic, and during cell division, Gef1 localizes to the division site. In all cases, cells were grown in YE5S to mid-log phase and imaged by spinning-disk confocal microscope in MatTek coverslip dishes at 25°C, under conditions that minimize any possible stress (Mutavchiev et al., 2016). In some cases, high exposures were used to emphasize absence of Gef1 from cell tips; therefore mitochondrial autofluorescence is apparent in images of Gef1-GFP and Gef1-3GFP. ***(B)*** Recruitment of Gef1-3YFP from cytosol to cell tips after treatment with the microtubule depolymerizing drug thiabendazole (TBZ; 150 μg/ml), which has off-target effects that lead to cell depolarization, independently of disrupting microtubules (Sawin and Snaith, 2004). TBZ was added just after imaging the zero time-point. After TBZ treatment, Gef1-3YFP transiently localizes to cell tips (red arrowheads) and later localizes more weakly to patches on cell sides (yellow arrowheads), which move towards cell middle. Scale bars, 10 μm. See also Video 4.

**Figure 6--figure supplement 1.**
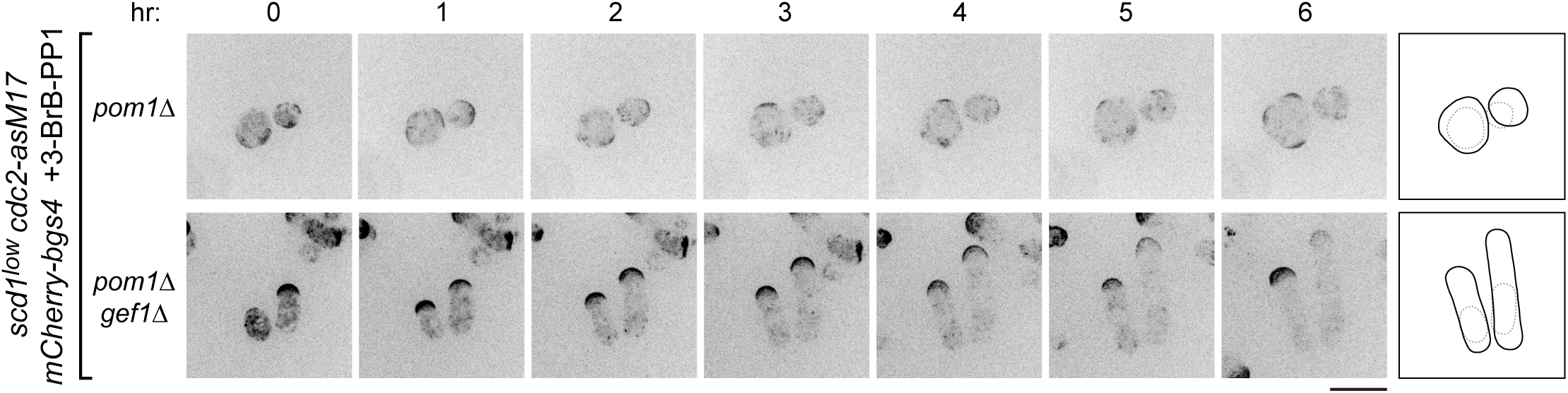
Deletion of *gef1* restores polarized growth to *scd1^low^ pom1Δ* cells. Time courses from movies showing mCherry-Bgs4 distribution and cell morphology in *pom1Δ* and *pom1Δ gef1Δ* cells, in *scd1^low^ cdc2-asM17 mCherry-bgs4* background, at the indicated times after start of imaging. *scd1* expression was repressed with thiamine for 24 hr before imaging. 3-BrB-PP1 was added 30 minutes before imaging. Diagrams show cell outlines at beginning and end of movies; outlines were aligned slightly to account for limited cell movement. Scale bar, 10 μm. See also Video 7.

**Figure 7--figure supplement 1.**
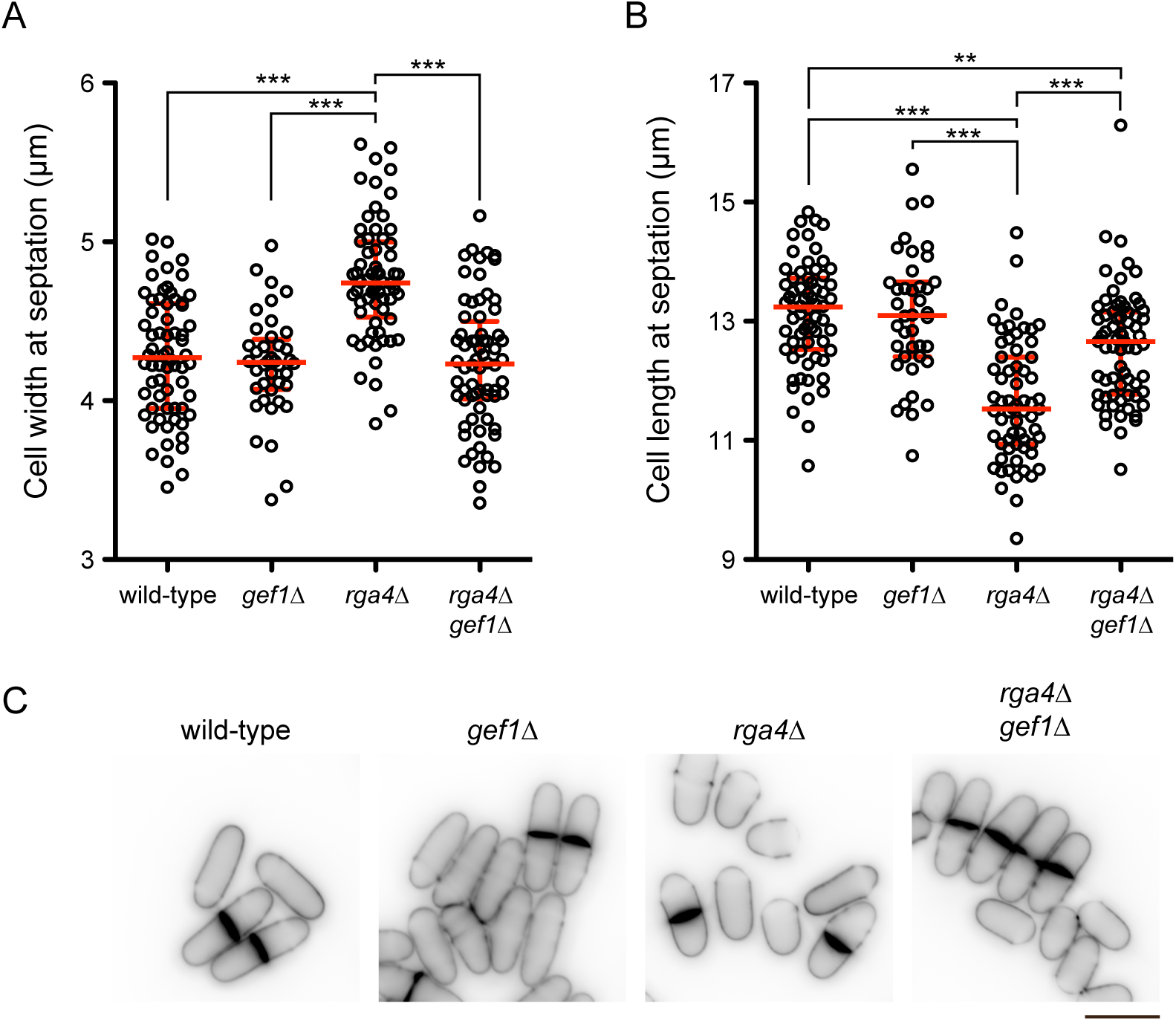
*gef1Δ* rescues the short/wide phenotype of *rga4Δ* cells. **(A)** Cell width at septation, and **(B)** cell length at septation, for cells of the indicated genotypes. Median and interquartile ranges are shown, with p-values for statistically significant differences (Student’s t-test, with Bonferroni correction); **, ***p*** = 0.001; ***, ***p*** < 0.0001. For each strain, 40-68 cells were scored. **(C)** Example images of calcofluor-stained cells used in measurements. Scale Bar, 10 μm.

